# PLEKHA4 Promotes Wnt/β-catenin Signaling-Mediated G1/S Transition and Proliferation in Melanoma

**DOI:** 10.1101/2020.07.29.225516

**Authors:** Adnan Shami Shah, Xiaofu Cao, Andrew C. White, Jeremy M. Baskin

**Affiliations:** Department of Chemistry and Chemical Biology, Cornell University, Ithaca, New York, 14853, USA; Weill Institute for Cell and Molecular Biology, Cornell University, Ithaca, New York, 14853, USA; Department of Biomedical Sciences, Cornell University, Ithaca, New York, 14853, USA

## Abstract

Melanoma patients incur substantial mortality, despite promising recent advances in targeted therapies and immunotherapies. In particular, inhibitors targeting BRAF-mutant melanoma can lead to resistance, and no targeted therapies exist for NRAS-mutant melanoma, motivating the search for additional therapeutic targets and vulnerable pathways. Here, we identify a regulator of Wnt/β-catenin signaling, PLEKHA4, as a factor required for melanoma proliferation and survival. PLEKHA4 knockdown in vitro leads to lower Dishevelled levels, attenuated Wnt/β-catenin signaling, and a block of progression through the G1/S cell cycle transition. In mouse xenograft models, inducible PLEKHA4 knockdown attenuated tumor growth in BRAF- and NRAS-mutant melanomas and synergized with the clinically used inhibitor encorafenib in a BRAF-mutant model. As an E3 ubiquitin ligase regulator with both lipid and protein binding partners, PLEKHA4 presents several opportunities for targeting with small molecules. Our work identifies PLEKHA4 as a promising drug target for melanoma and clarifies a controversial role for Wnt/β-catenin signaling in the control of melanoma proliferation.

## INTRODUCTION

Melanoma is the most aggressive and deadliest form of skin cancer. The root cause of most melanomas is somatic mutations in a relatively small number of genes (1). Roughly 65% of melanoma cases feature a V600D/E mutation in the Ser/Thr kinase BRAF, and an additional 10% feature a Q61K/R mutation in the GTPase NRAS (2). These genetic alterations cause phenotypic changes, including elevated signaling through MAP kinase, PI 3-kinase, and other related pathways, which lead to increased cell proliferation, differentiation, and ultimately tumorigenesis and malignancy (3).

Inhibitors of BRAF or the downstream kinase MEK heralded an era of targeted therapies for BRAF-mutant melanomas (4–8). Nonetheless, resistance typically occurs in roughly one year, leading to relapse, and no targeted therapies exist for NRAS-mutant melanomas (9–11). Further, immunotherapies, such as checkpoint inhibitors, have more long-lasting effects but are only successful in a subset of patients (12,13). Combinations of BRAF targeted therapies and anti-PD1 immunotherapies are promising avenues but are still not universally effective (14). Thus, new therapeutic strategies are needed to prevent melanomagenesis and progression.

Wnt/β-catenin signaling, which regulates proliferation, is aberrantly hyperactive in several cancers, including melanoma (15). In the canonical, β-catenin-dependent form of this pathway, secreted Wnt ligands engage a receptor from the Frizzled family in the plasma membrane of the Wnt-receiving cell (16,17). This binding event causes recruitment of Dishevelled (DVL), which mediates disassembly of a multicomponent β-catenin destruction complex, resulting in β-catenin stabilization, nuclear translocation, and altered gene expression at several loci, most notably those associated with the TCF/LEF transcription factor family. In cancer, aberrant Wnt/β-catenin signaling leads to increased expression of Wnt/β-catenin target genes including Cyclin D1 and c-Myc, which regulate progression through the G1/S transition of the cell cycle, helping to promote proliferation, tumorigenesis, and malignancy (15).

Wnt signaling pathways have been linked to melanoma, but their exact roles remains controversial (15,18–20). Wnt/β-catenin signaling has been shown to promote melanoma tumor initiation and growth in both BRAF and NRAS mutant backgrounds (21–24). Further, a recent study using a new engineered mouse model implicated Wnt signaling in the transformation of healthy melanocyte stem cells to melanoma in a BRAF and PTEN mutant background (25). As well, BRAF inhibition is more effective in settings with lower levels of β-catenin (26). Yet, elevated levels of nuclear (active) β-catenin have correlated with diverging patient survival, depending on the study (19,27–30). Beyond the controversial roles of Wnt/β-catenin signaling in melanoma, β-catenin-independent non-canonical Wnt signaling controls actin cytoskeletal dynamics and cell migration and has been implicated in melanoma metastasis (31,32). In fact, melanoma progression has been proposed to involve a phenotype switching model wherein the canonical and non-canonical pathways alternate to allow cells to switch between a proliferative and migratory phenotypes (15,33). Thus, Wnt signaling pathways appear to be important players in melanoma progression in most contexts and are thus a potential point of therapeutic intervention.

Numerous efforts have been made to drug Wnt signaling in cancer (30,34,35). These efforts have largely focused on inhibiting core Wnt components (e.g., PORCN, FZD, β-catenin/CBP) (36). Though efficacious in model systems, they have seen limited success in vivo due to undesirable side effects on homeostatic Wnt signaling in non-diseased tissues (37). Fortunately, Wnt/β-catenin signaling is subject to many levels of regulation, and though core Wnt components are typically essential due to important roles in development and tissue homeostasis, many modulators, or tuners, of Wnt signaling strength may not be required for viability (16,17,34). Thus, it is a high priority to identify modulators of Wnt signaling, whose inhibition downregulates but does not completely eliminate Wnt signaling, as potential therapeutic targets.

Among the many factors involved in Wnt signaling, DVL has emerged as a major point of regulation (38,39). Several different E3 ubiquitin ligases act on DVL, modulating its levels and thus changing the strength of the Wnt signal in Wnt-receiving cells (40–46). To this end, we recently discovered that the phosphoinositide-binding protein PLEKHA4 (pleckstrin homology containing family A, number 4) modulates the activity of the CUL3– KLHL12 E3 ligase that polyubiquitinates DVL (40,47). PLEKHA4 acts to sequester the substrate-specific adaptor KLHL12 within plasma membrane-associated clusters, thus reducing DVL ubiquitination, increasing DVL levels, and enhancing Wnt/β-catenin signaling in mammalian cells. Thus, PLEKHA4 acts as a tuner for DVL levels and Wnt signaling strength, as near-complete elimination of PLEKHA4 resulted in only partial DVL depletion and attenuation of Wnt signaling.

Intriguingly, among >20 cancers investigated in a TCGA dataset, PLEKHA4 expression was highest in melanoma (**Figure 1A**); however, its levels are low in healthy melanocytes as analyzed in the Genevestigator database (2,48). We were thus motivated to test whether PLEKHA4 is an important factor for promoting pathological Wnt signaling in melanoma, as a step toward both validating Wnt/β-catenin signaling in general, and PLEKHA4 in particular, as therapeutic targets in melanoma. Here, we report that melanoma cells from both BRAF and NRAS mutant backgrounds require PLEKHA4 for survival and proliferation in vitro and in vivo in mouse xenograft and allograft models. Depletion of PLEKHA4 by siRNA and shRNA led to attenuated Wnt signaling in these models and phenocopied inhibitors or siRNA knockdown of core Wnt components. Further, inducible PLEKHA4 knockdown in the presence of the clinically used BRAF V600D/E inhibitor encorafenib (49) displayed a striking synergistic effect in a xenograft model of BRAF-mutant melanoma, suggesting the therapeutic potential of targeting PLEKHA4 in melanoma. This work highlights PLEKHA4 as a new modulator of Wnt/β-catenin signaling strength in melanoma that, by promoting G1/S cell cycle transition, maintains cell proliferation in melanoma. Importantly, our study provides additional clarity on the pathological role of Wnt/β-catenin signaling in this disease and suggests that pharmacological inhibition of PLEKHA4 could represent a promising new avenue for targeted therapy in melanoma.

**Figure 1.**
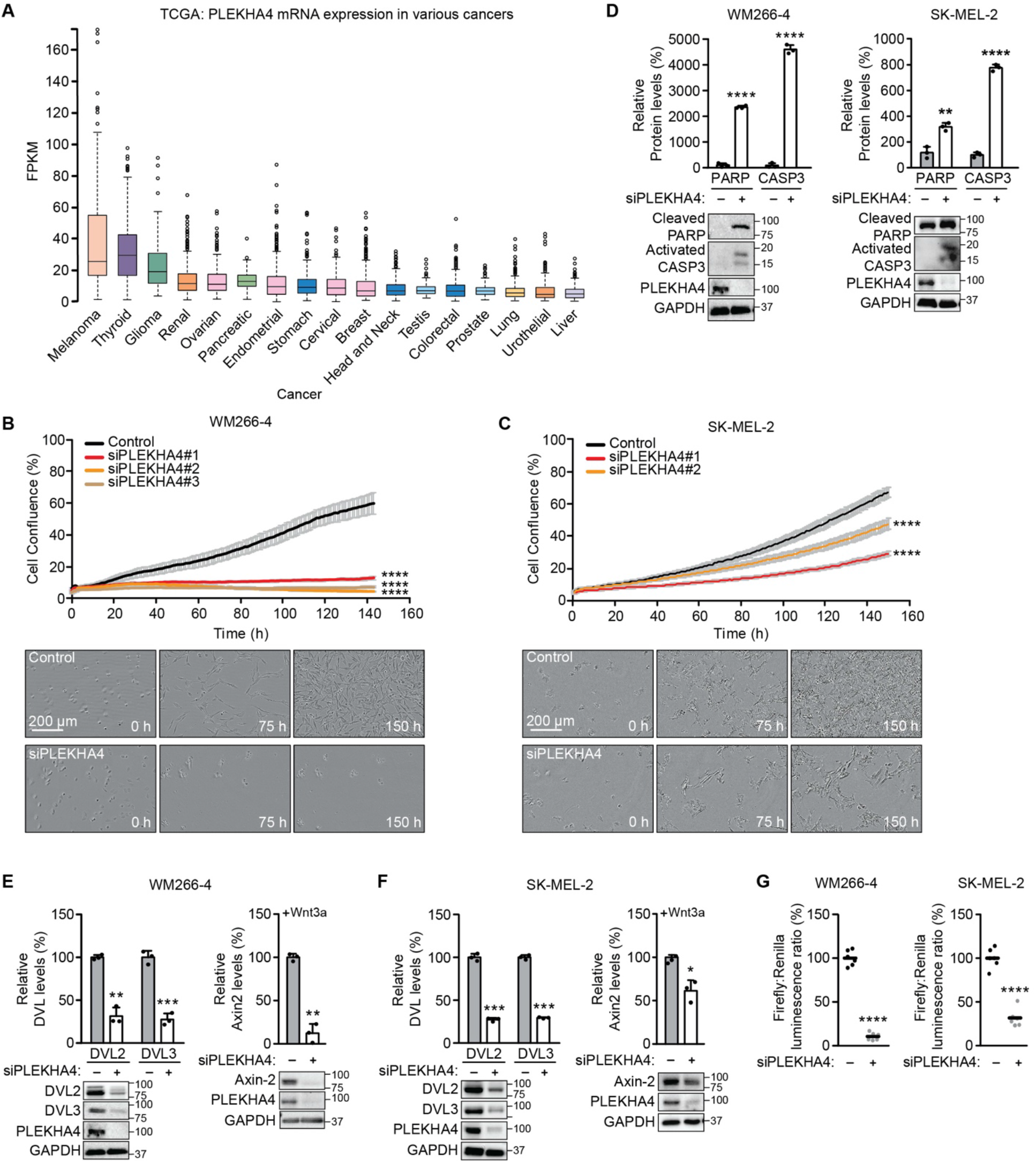
PLEKHA4 loss from melanoma cells reduces proliferation and increases apoptosis via attenuation of Wnt/β-catenin signaling. (A) Analysis of PLEKHA4 mRNA levels in various cancers, based on data generated by the TCGA Research Network. FPKM, fragments per kilobase of transcript per million mapped reads. (B and C) PLEKHA4 knockdown by siRNA inhibits melanoma cell proliferation in vitro. Automated brightfield imaging of cell proliferation via IncuCyte of (B) WM266-4 and (C) SK-MEL-2 melanoma cells treated with siRNA duplexes targeting different regions of PLEKHA4 (siPLEKHA4 #1, #2 and #3) or a negative control siRNA (n=3). (D–F) PLEKHA4 knockdown causes increased levels of apoptotic markers (cleaved PARP and activated Caspase 3 (CASP3)) and reduction in Wnt signaling (DVL2, DVL3, and Axin2) in mutant melanoma cells. Shown is Western blot analysis of WM266-4 and SK-MEL-2 cells subjected to siPLEKHA4 or a negative control siRNA (−) (n=3). For Axin2 analysis, cells were stimulated with Wnt3a-containing conditioned media concurrently with siRNA. (G) PLEKHA4 modulates Wnt/β-catenin signaling in WM266-4 and SK-MEL-2 cells. Shown is TOPFlash assay signal, i.e., ratio of β-catenin-dependent firefly luciferase activity to constitutive Renilla luciferase activity in WM266-4 or SK-MEL2 cells treated with siPLEKHA4 or negative control siRNA (−) and stimulated with Wnt3a-containing conditioned media (n=6). * p < 0.05, ** p < 0.01, *** p<0.001, **** p < 0.0001. Scale bars: 200 μm.

## RESULTS

### PLEKHA4 knockdown blocks proliferation and increases apoptosis in melanoma cells

In the course of earlier work on PLEKHA4 in HeLa cells (47), we noticed that, qualitatively, its knockdown by siRNA had mild effects on cell proliferation and viability. We reasoned that cancer cells expressing highest levels of PLEKHA4 might be more sensitive to its loss. Analysis of patient gene expression data in the TCGA database revealed widespread expression of PLEKHA4 in many types of cancers (**Figure 1A**) (2). Yet, its levels were highest in melanoma cancer cells, and intriguingly, its levels in healthy melanocytes, as analyzed in the Genevestigator database, were low (48). With a working hypothesis that PLEKHA4 might be an important factor in melanomagenesis and progression, we examined its requirement for proliferation and survival in two melanoma cell lines: WM266-4, a BRAFV600D mutant line, and SK-MEL-2, an NRAS Q61R mutant line.

We validated several PLEKHA4 siRNA duplexes (**Figure S1A**) and examined effects of PLEKHA4 knockdown using automated, continual monitoring of cell number on IncuCyte system, wherein images were acquired every hour for 150 h. We observed a strong reduction of cell proliferation upon PLEKHA4 knockdown in both cell lines, using multiple siRNA duplexes (**Figure 1B–C**). Examination of the images suggested substantial cell death was occurring, and indeed, Western blot analysis of lysates from these cells revealed that PLEKHA4 knockdown caused increases in levels of cleaved PARP and activated caspase 3, two markers of apoptosis (**Figure 1D**).

### PLEKHA4 promotes Wnt/β-catenin signaling in melanoma cells

Given the role of PLEKHA4 as a positive regulator of Wnt/β-catenin signaling in other cells (47), we next investigated effects of PLEKHA4 knockdown on Wnt signaling in the context of melanoma. We found that siRNA-mediated PLEKHA4 knockdown led to reduced levels of DVL2 and DVL3, the two major DVL isoforms in both the BRAF and NRAS mutant melanoma cell lines (**Figure 1E–F**). We then examined effects on Wnt/β-catenin signaling using two approaches. First, PLEKHA4 knockdown led to a >50% decrease in luminescence from melanoma cells stably expressing a β-catenin-dependent luciferase transcriptional reporter (TOPFlash) that were stimulated with Wnt3a (**Figure 1G**). Second, we found that PLEKHA4 knockdown in cells stimulated with Wnt3a led to reduced levels of Axin2, whose expression is induced by canonical Wnt/β-catenin signaling, by Western blot (**Figure 1E–F**).

To complement these studies on PLEKHA4 knockdown, we examined whether perturbing Wnt signaling via two distinct mechanisms would similarly affect viability and proliferation of these melanoma cells. First, we used a pan Wnt inhibitor (IWP-4) that targets Porcupine, an O-acyltransferase that installs a posttranslational modification that is required for their secretion from Wnt-producing cells and thus for Wnt signaling (50). We found that both IWP-4 treatment led to a drastic cell proliferation defect upon IWP-4 treatment in both the cell lines (**Figures 2A** and **S1B**). Second, we performed siRNA treatment of DVL2 or DVL3, the direct mechanistic targets of PLEKHA4 action (47), and found a similar effect on cell proliferation in both melanoma cell lines (**Figures 2B** and **S1C**). Further, Western blot analyses on DVL2 or DVL3 knockdown samples revealed increases in the levels of cleaved PARP and activated caspase 3, suggesting increases in apoptosis similar to PLEKHA4 knockdown (**Figure 2C**). Together, these data indicate that PLEKHA4 acts as a positive modulator of Wnt/β-catenin signaling in melanoma and suggests that it mediates cell survival and proliferation in melanoma.

**Figure 2.**
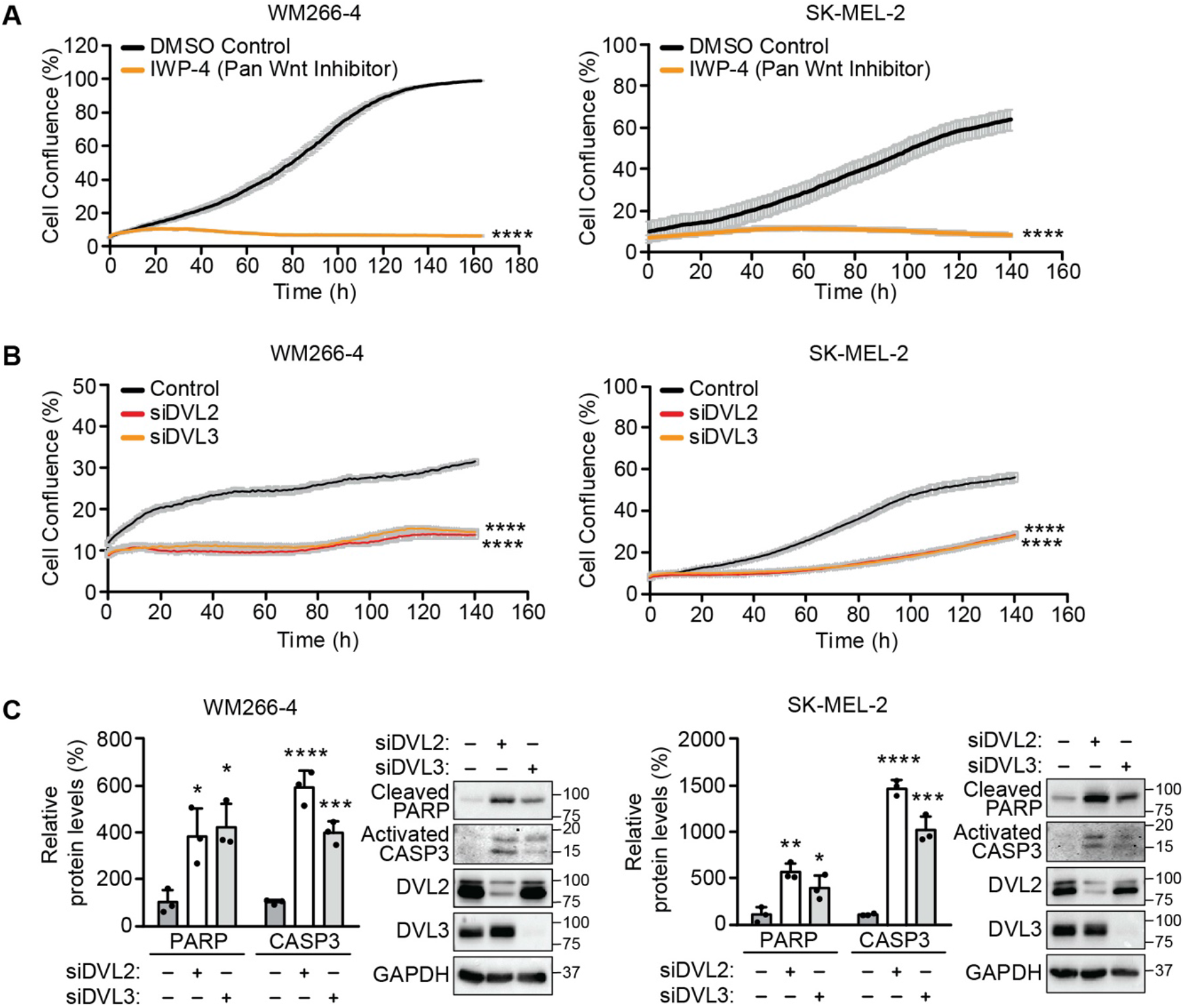
Inhibition of Wnt signaling decreases proliferation and induces apoptosis in melanoma cells. (A and B) Pan Wnt inhibition via treatment with the inhibitor IWP-4, compared to DMSO control (A), and knockdown of DVL2 (siDVL2) or DVL3 (siDVL3), compared to negative control siRNA (B), both attenuate proliferation of WM266-4 and SK-MEL-2 cells, as assessed by automated brightfield imaging of cell proliferation using an IncuCyte system (n=3). (C) Knockdown of DVL2 or DVL3 causes apoptosis in WM266-4 and SK-MEL-2 cells. Shown is Western blot analysis of WM266-4 and SK-MEL-2 cells subjected to siDVL2, siDVL3, or a negative control siRNA (−) (n=3). * p < 0.05, ** p < 0.01, *** p<0.001, **** p < 0.0001.

### PLEKHA4 regulates the G1/S transition and melanoma cell proliferation

A major role of Wnt/β-catenin signaling is to stimulate proliferation by promoting progression through the G1/S cell cycle transition. The effects of PLEKHA4 and Wnt perturbation on cell growth curves suggested an effect on proliferation, and we next examined whether the mechanism of action of PLEKHA4 occurred via perturbing the cell cycle. First, we analyzed cell cycle phase on asynchronous WM266-4 cells treated with either control or two different PLEKHA4 siRNA duplexes and stained fixed cells with propidium iodide to measure DNA content. We found that PLEKHA4 knockdown led to an accumulation of cells in the G1 phase (**Figure 3A**). Importantly, this PLEKHA4 knockdown-induced G1/S transition defect could be rescued by introduction of an siRNA-resistant form of PLEKHA4 via lentiviral transduction (**Figures 3D** and **S2**). Intriguingly, effects of PLEKHA4 knockdown could also be rescued by overexpression of DVL2 or DVL3, the downstream targets of PLEKHA4, strongly suggesting that the established mechanism of action of PLEKHA4 on DVL proteins, via effects on their ubiquitination by CUL3–KLHL12 (47), occurs as well in these melanoma cells.

**Figure 3.**
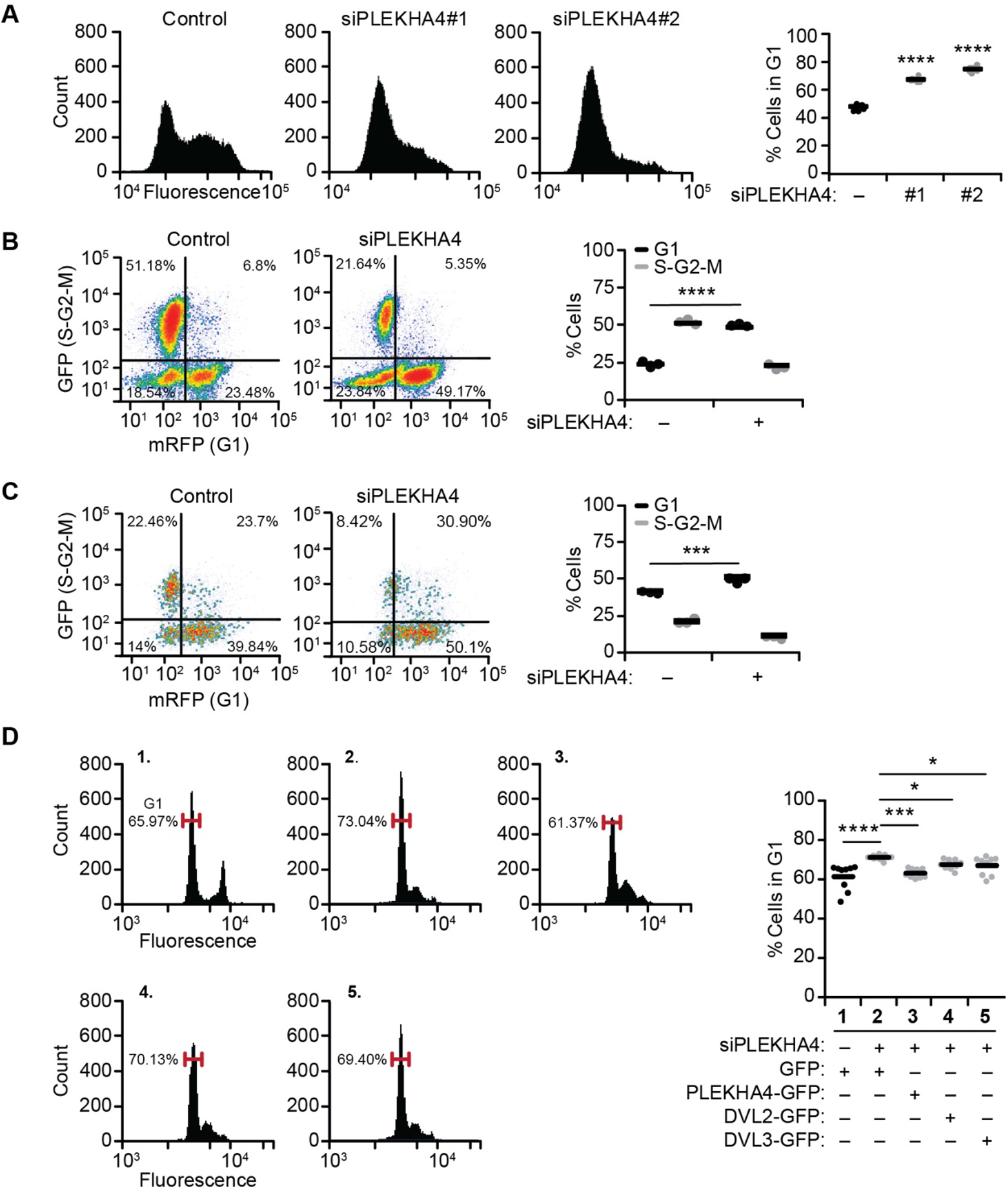
PLEKHA4 knockdown inhibits Wnt/β-catenin mediated G1/S cell cycle transition. (A) PLEKHA4 knockdown leads to accumulation of WM266-4 cells in G1 phase. An asynchronous population of WM266-4 cells was treated with one of two different siRNA duplexes against PLEKHA4 (siPLEKHA4 #1 and #2) or a negative control siRNA (−), followed by fixation, propidium iodide staining, and flow cytometry analysis. (n=6) (B and C) PLEKHA4 knockdown impairs G1/S transition in synchronized melanoma cells stably expressing the FUCCI cell cycle indicator. WM266-4-FUCCI (B) and SK-MEL-2-FUCCI (C) stable cells were synchronized to G1 phase via serum starvation and concurrent treatment with the indicated siRNA duplex for 48 h. Cells were then released into fresh medium containing FBS, followed by the quantification of mRFP (G1) and GFP (S-G2-M) fluorescence via flow cytometry (n=3). (D) PLEKHA4-GFP, DVL2-GFP, and DVL3-GFP can rescue the attenuation of the G1/S transition defect induced by PLKEHA4 knockdown. WM266-4 cells were synchronized to G1 phase, subjected to siPLEKHA4 or negative control siRNA (−), and stimulated with media containing FBS and simultaneously transduced with conditioned media containing lentivirus encoding GFP, siRNA-resistant PLEKHA4-GFP, DVL2-GFP, or DVL3-GFP, followed by fixation, propidium iodide staining, and flow cytometry analysis (n=9). * p < 0.05, *** p<0.001, **** p < 0.0001.

To examine the G1/S phenotype in more detail, including its dynamics, we used the FUCCI (Fluorescent Ubiquitination-based Cell Cycle Indicator) system, a live cell-compatible, dual color reporter that wherein cells in G1 phase express mRFP (red) and cells in S, G2, or M phase express GFP (green). We generated WM266-4 and SK-MEL-2 cell lines stably expressing the FUCCI probes and synchronized either control or PLEKHA4 knockdown cells to G1 using serum starvation (51). Upon release from this G1 arrest by addition of serum, we found that, for both cell lines, PLEKHA4 knockdown caused an increase in retention in G1 phase, i.e., a failure to progress to S phase (**Figure 3B–C**).

To complement this phenotypic characterization of G1/S defects, we examined levels of Cyclin D1 and c-Myc, two well-studied transcriptional targets of Wnt/β-catenin signaling that affect the G1/S cell cycle transition (52,53). In asynchronous populations of WM266-4 or SK-MEL-2 cells, we found that PLEKHA4 knockdown led to decreased levels of both Cyclin D1 and Myc (**Figure 4A–B**). Further, PLEKHA4 knockdown on G1-synchronized WM266-4 or SK-MEL-2 cells (via serum starvation) led to a similar decrease in the levels of Cyclin D1 and Myc, as well as DVL2 and DVL3 (**Figure 4C–D**). Importantly, a decrease in the levels of these proteins induced by PLEKHA4 knockdown could be rescued by lentiviral transduction with the siRNA-resistant form of PLEKHA4 (**Figure 4E**). Interestingly, the decrease in levels of Cyclin D1 and Myc induced by PLEKHA4 knockdown could also be rescued by expression of DVL2 or DVL3, further supporting the proposed mechanism of action (**Figure 4E**). Finally, to complement these findings, we found that DVL2 or DVL3 knockdown led to the same effects on Cyclin D1 and Myc levels in both melanoma cell lines (**Figure 4F**). Overall, these data indicate that decreasing PLEKHA4 levels in melanoma leads to a Wnt/β-catenin-mediated G1/S cell cycle transition defect via effects on the key proliferation markers Cyclin D1 and Myc.

**Figure 4.**
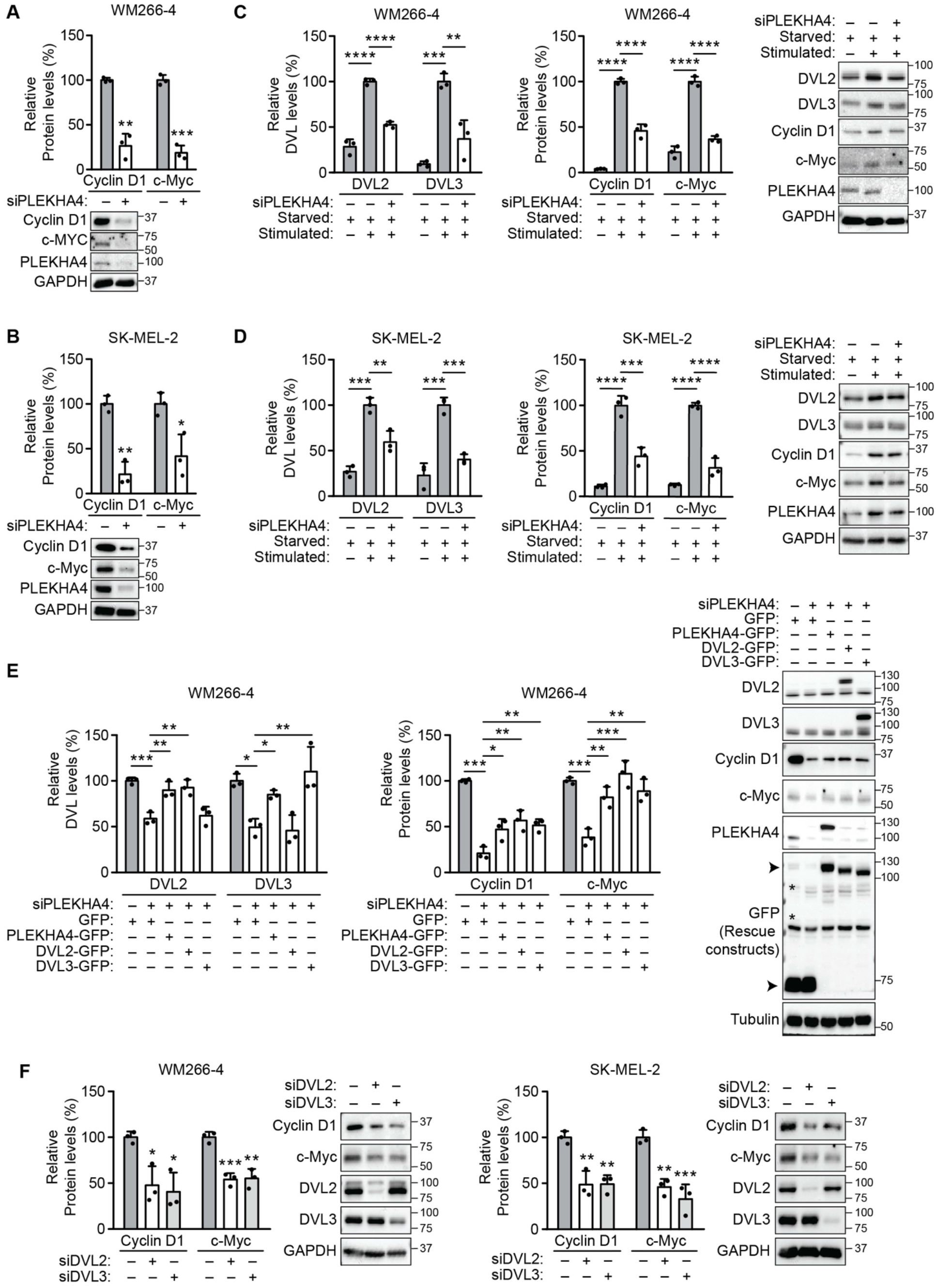
PLEKHA4 knockdown reduces levels of Wnt/β-catenin-controlled markers of proliferation. (A and B) PLEKHA4 knockdown decreases Cyclin D1 and c-Myc levels in asynchronous WM266-4 (A) and SK-MEL-2 (B) cells. Shown is Western blot analysis and quantification of lysates from the indicated cells treated with an siRNA duplex against PLEKHA4 (siPLEKHA4) or a negative control siRNA (−) (n=3). (C and D) PLEKHA4 modulates the levels of DVL2, DVL3, Cyclin D1, and c-Myc in G1-synchronized WM266-4 (C) and SK-MEL-2 (D) cells. Shown is Western blot analysis and quantification of lysates from melanoma cells synchronized to G1 phase via serum starvation that were treated with siPLEKHA4 or a negative control siRNA (−) and then stimulated with FBS-containing medium (n=3). (E) PLEKHA4-GFP, DVL2-GFP, and DVL3-GFP can rescue the changes in DVL2, DVL3, Cyclin D1, and c-Myc levels induced by PLEKHA4 knockdown in WM266-4 cells. Shown is Western blot analysis and quantification of lysates from WM266-4 cells subjected to siPLEKHA4 or negative control siRNA (−) and transduced with conditioned media containing lentivirus encoding GFP, siRNA-resistant PLEKHA4-GFP, DVL2-GFP, or DVL3-GFP (n=3). (F) Knockdown of DVL2 or DVL3 leads to a decrease in levels of Cyclin D1 and c-Myc. Shown is Western blot analysis and quantification of lysates from WM266-4 and SK-MEL-2 cells treated with the indicated siRNA duplex or negative control siRNA (n=3). * p < 0.05, ** p < 0.01, *** p<0.001, **** p < 0.0001.

### PLEKHA4 is required for tumorigenic and malignant properties in melanoma in vitro

The above molecular and phenotypic data implicate PLEKHA4 as a novel modulator of Wnt signaling in melanoma whose attenuation causes defects in cell cycle progression and proliferation. We therefore envisioned that loss of PLEKHA4 in melanoma cells might attenuate cancer-causing properties in vitro such as clonogenic capacity, or the ability of a single cell to proliferate into a colony.

We first examined effects of PLEKHA4 knockdown on the anchorage-dependent clonogenic capacity of melanoma cells, using crystal violet staining of colonies derived from single cells grown on traditional 2D cell culture surfaces. PLEKHA4 knockdown in both WM266-4 and SK-MEL-2 cell lines led to substantial losses in clonogenic capacity: 80% for the BRAF-mutant WM266-4 and 50% for the NRAS-mutant SK-MEL-2 (**Figure 5A**). Further, a similar effect was observed upon inhibition of Wnt signaling via other mechanisms, including IWP-4 treatment (**Figure 5E**) and knockdown of DVL2 or DVL3 (**Figure 5C**).

**Figure 5.**
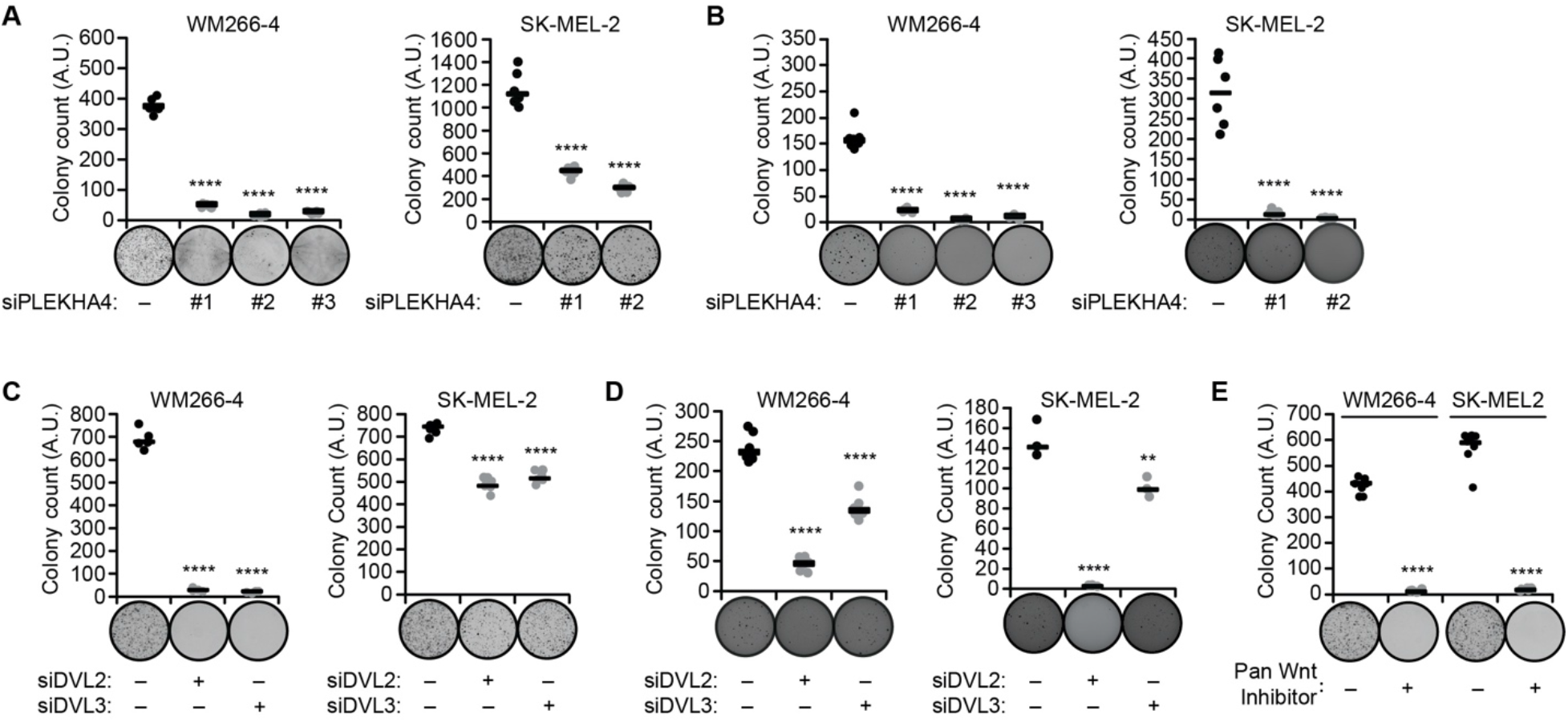
PLEKHA4 knockdown and Wnt inhibition causes loss of tumorigenic and malignant properties in melanoma cells in vitro. Cells treated as described below were analyzed via anchorage-dependent colony formation assay with crystal violet staining (A, C, and E) or anchorage-independent soft agar assay (B and D). Representative images are shown for each treatment, and graphs indicate colony count. (A and B) Cells were treated with the indicated siRNA duplex against PLEKHA4 or negative control siRNA (−) (n=6). (C and D) Cells were treated with siRNA duplexes against DVL2, DVL3, or negative control siRNA (n=6 for all, except for n=3 for SK-MEL-2 in (D)). (E) Cells were treated with the pan Wnt inhibitor IWP-4 or DMSO control (−) (n=6). ** p < 0.01, **** p < 0.0001.

To evaluate tumorigenic potential of malignant cells grown in a soft substrate that better mimics the tumor environment, we employed an anchorage-independent colony formation assay (54). Here, colony formation was measured after seeding cells in a 3D soft agar environment, followed by nitrotetrazolium blue staining. We found that PLEKHA4 knockdown in both melanoma cell lines strongly reduced anchorage-independent growth capacities (**Figure 5B**). Again, inhibition of Wnt signaling via DVL2 or DVL3 knockdown led to a similar decrease in anchorage-independent growth (**Figure 5D**). These data indicate that loss of PLEKHA4 causes a drastic decrease in tumorigenic and malignant properties in BRAF and NRAS mutant melanoma in vitro.

### PLEKHA4 knockdown attenuates melanoma tumor growth in vivo

Buoyed by the in vitro data implicating PLEKHA4 as a factor required for melanoma cell proliferation, we next tested whether PLEKHA4 played a similar role in vivo. Here, we used two different types of mouse models. First, we established xenografts in immunocompromised NOD *scid* gamma (NSG) mice using WM266-4 and SK-MEL-2 cells, the BRAF- and NRAS-mutant human melanoma cell lines that we had used for the extensive in vitro studies above. Separately, to assess effects of PLEKHA4 knockdown within wild-type mice, we established allografts in C57BL6.J mice using the YUMM1.7 cells, a syngeneic engineered mouse melanoma cell line bearing several mutations commonly found within melanoma, including BRAF V600E mutation, as well as mutations in PTEN and CDKN2A (55).

For these in vivo experiments, we established PLEKHA4 knockdown by generating cell lines stably expressing a doxycycline-inducible shRNA against human and mouse PLEKHA4. To accomplish this, we generated stable cell lines expressing several different shRNA constructs against human PLEKHA4 in WM266-4 cells (**Figure S3A**) and against mouse PLEKHA4 in YUMM1.7 cells (**Figure S4**). Cells were grown in vitro, and PLEKHA4 knockdown was induced by addition of doxycycline, and Western blot analysis was performed. We examined PLEKHA4 levels and additionally levels of DVL2, DVL3, Cyclin D1, and Myc to determine the most effective shRNAs from each collection (**Figures S3A** and **S4**). We further validated the effectiveness of these shRNAs at suppressing Wnt3a-stimulated Wnt/β-catenin signaling using the TOPFlash system within these PLEKHA4 stable knockdown lines (**Figure S3B**). The three best-performing shRNAs against human PLEKHA4, as validated in WM266-4 cells, were subsequently stably expressed and validated in SK-MEL-2 cells (**Figure S5**).

We then generated xenograft/allograft models by subcutaneous injection into the shoulder or hind leg flanks in the absence of doxycycline to allow tumors to form. For the WM266-4 xenograft and YUMM1.7 allografts, after 12 days in the absence of doxycycline to allow tumors to form, doxycycline was administered for 10–12 days to induce PLEKHA4 knockdown (**Figure 6A**). As negative controls, stable cell lines expressing luciferase shRNA were employed. We monitored tumor progression over this time span and observed a major attenuation of tumor growth for both of these BRAF-mutant models (**Figure 6B–C**). Further analysis of the tumors at the experimental endpoint revealed that PLEKHA4 tumors were approximately four-fold smaller in the WM266-4/NSG model (**Figure 6B**) and three-fold smaller in the YUMM1.7/C57BL6.J model (**Figure 6C**).

**Figure 6.**
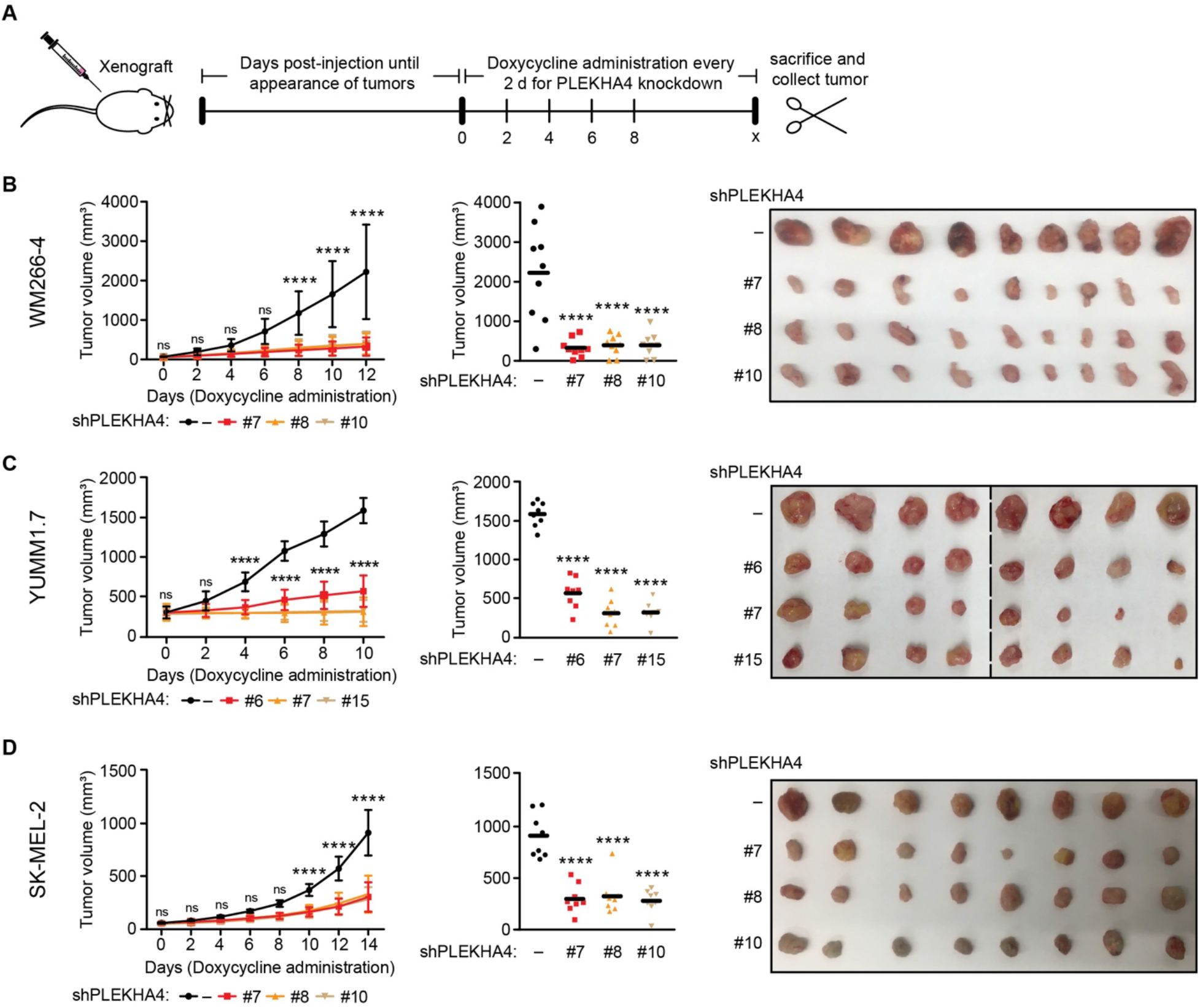
Inducible PLEKHA4 knockdown inhibits melanoma tumor xenograft/allograft growth *in vivo*. (A) Schematic representation of experimental setup and timeline for xenograft/allograft analyses. Cell lines stably expressing doxycycline-inducible shRNA against human (WM266-4 and SK-MEL-2) or mouse (YUMM1.7) PLEKHA4 or a negative control shRNA (−) were xenografted into NSG (WM266-4 and SK-MEL-2) and C57BL/6J (YUMM1.7) mice. Mice were monitored, and after small tumor bumps appeared (12 d for WM266-4 and YUMM1.7; 45 d for SK-MEL-2), doxycycline was administered through the drinking water for a total of 10–16 d to induce PLEKHA4 knockdown. Tumor progression over this time period was monitored by measurement of tumor dimensions using a digital caliper and calculation of tumor volume using the formula v = 0.5233*l*w^2^. Mice were then sacrificed, and tumors were collected (n=12 for WM266-4-xenografted NSG mice, n=10 for YUMM1.7-allografted C57BL/6J mice, and n=14 for SK-MEL-2-xenografted NSG mice). (B–D) Data from studies using WM266-4 xenografts (B), YUMM1.7 allografts (C), and SK-MEL-2 xenografts (D). The plots at left show changes in tumor volume over time, and the plot in the middle show final tumor volumes measured with a caliper post-harvesting, with images of tumors harvested at the endpoint shown at right. n=9 for WM266-4 and n=8 for YUMM1.7 and SK-MEL-2. ns, not significant, **** p < 0.0001.

To test the effect of PLEKHA4 knockdown on NRAS-mutant melanoma in vivo, we established an SK-MEL-2 xenograft, and once visible tumors appeared at 1.5 months post-injection, doxycycline administration was carried out for 14 days (**Figure 6A**). Analysis of tumor progression and endpoint data revealed that tumor growth was attenuated two-fold in the PLEKHA4 knockdown samples compared to control (**Figure 6D**). These data demonstrate that PLEKHA4 knockdown in an in vivo, tumor xenograft or allograft setting results in a substantial decrease in tumor growth and implicate PLEKHA4 and, by extension, Wnt signaling, as a regulator of BRAF and NRAS-mutant melanoma progression in vivo.

### PLEKHA4 knockdown synergizes with a BRAF inhibitor in vivo

Finally, we wanted to establish the feasibility of targeting PLEKHA4 in a model of a therapeutic setting. Targeted BRAF therapy, i.e., BRAF or MEK inhibitors, represents a frontline treatment for melanoma (4,6). Though effective, this treatment has its limitations, including resistance, leading to relapse (10,11). PLEKHA4 and its effect on Wnt signaling could represent a second, parallel druggable pathway to block melanoma progression. Thus, we examined whether PLEKHA4 knockdown could synergize with treatment with encorafenib (BRAFi), an FDA-approved BRAF inhibitor used routinely to treat BRAF-mutant melanoma (49). We generated WM266-4 xenografts bearing doxycycline-inducible PLEKHA4 or control shRNA as before. On day 12 post-injection, following the formation of tumors, mice were administered both doxycycline to induce shRNA expression and encorafenib, via daily oral gavage, to inhibit BRAF and downstream MAP kinase signaling (**Figure 7A**).

**Figure 7.**
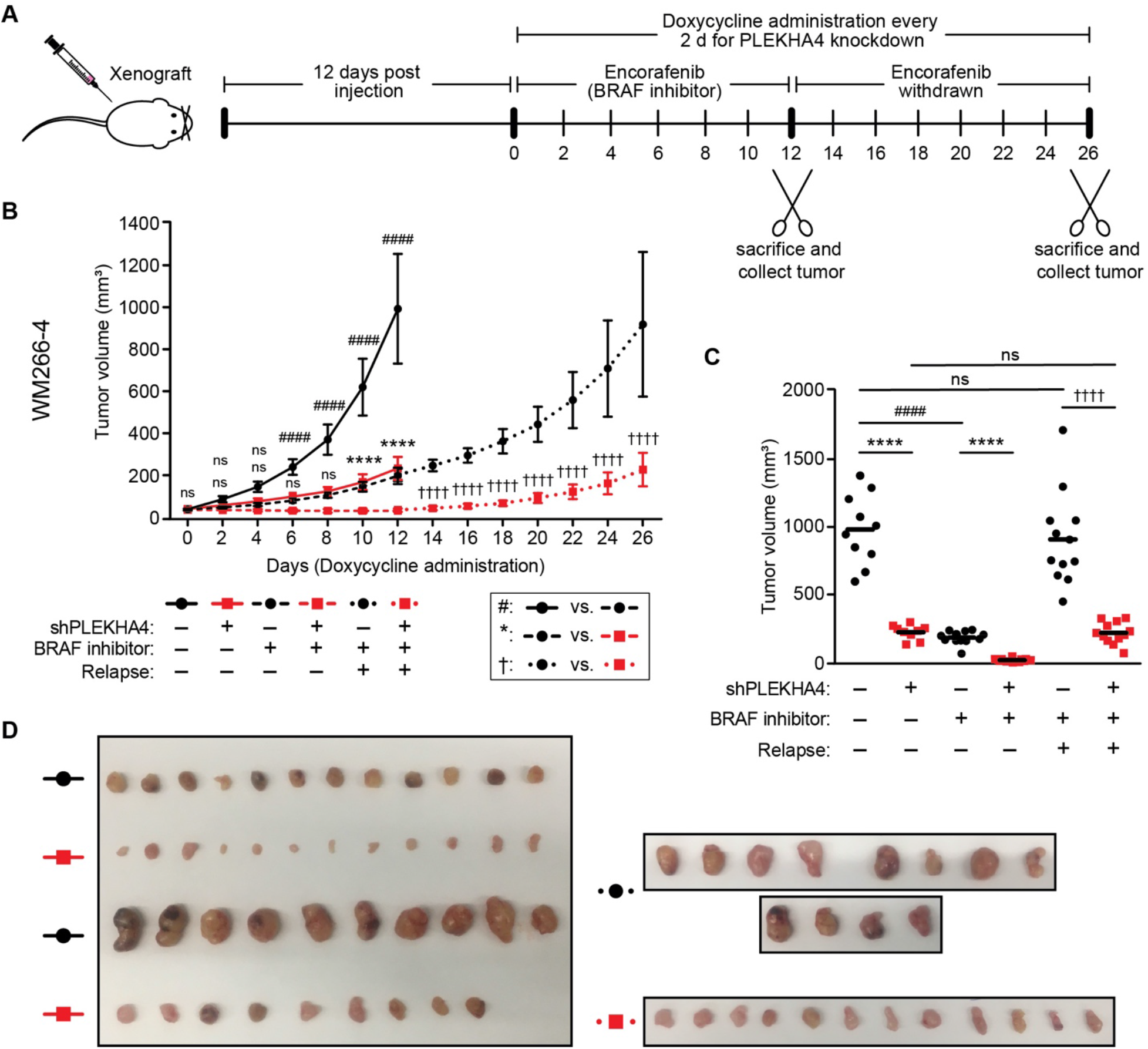
PLEKHA4 knockdown synergizes with the BRAF inhibitor encorafenib to attenuate melanoma tumor xenograft growth in vivo. (A) Schematic representation of experimental setup and timeline. WM266-4 cells stably expressing a doxycycline-inducible shRNA hairpin against PLEKHA4 (shPLEKHA4 #7) or a control shRNA (−) were xenografted into NSG mice. Mice were monitored, and after tumors became visible 12 d post-injection (labeled as day 0), doxycycline was administered through the drinking water and the BRAF inhibitor encorafenib or vehicle control was administered via oral gavage every day for 12 d. On day 12, all vehicle-treated mice and half of the encorafenib-treated mice bearing control and PLEKHA4 knockdown tumors were sacrificed for tumor collection. For the remaining mice, doxycycline treatment was continued but encorafenib was withdrawn to assess effect of PLEKHA4 knockdown on relapse for another 14 d. On day 26, mice were sacrificed for tumor collection. (B–D) Data from these studies. (B) Plot showing changes in tumor volume over time, with dimensions determined as described in the Figure 6 legend. (C) Final tumor volumes measured with a caliper post-harvesting. (D) Images of tumors harvested at endpoints: day 12 (left) and day 26 (right) (n=10–12 for each group). ns, not significant, ****, ####, and ††††: p < 0.0001.

This study was divided into two phases. In the first phase, we examined effects of PLEKHA4 knockdown and encorafenib treatment separately or in combination. We found that encorafenib treatment prevented tumor growth compared to control, similar to effects of PLEKHA4 knockdown alone (**Figure 7B**). Encouragingly, encorafenib treatment in the context of PLEKHA4 knockdown resulted in significantly reduced tumor growth compared to either PLEKHA4 knockdown or encorafenib treatment alone (**Figure 7B**). These data suggest a strong synergy between PLEKHA4 knockdown and BRAF inhibition. Further analysis of tumor size and endpoint data confirmed that encorafenib treatment and PLEKHA4 knockdown exhibited similar effects on tumor size compared to control samples (**Figure 7C–D**).

Clinically, melanoma tumors can relapse upon development of resistance to BRAF inhibitors such as encorafenib, as well as withdrawal of the inhibitor (10,11). This relapse is problematic, leading to further disease progression and poor patient outcomes. We were thus motivated to examine the effects on tumor growth of continued PLEKHA4 inhibition after removal of encorafenib. In the second phase of the study, we extended the study on both control and PLEKHA4 knockdown groups that had been treated with encorafenib during the first phase of the study. Here, we removed encorafenib but continued doxycycline treatment for an additional 14 days to sustain PLEKHA4 knockdown in a setting that models relapse following a BRAF inhibitor. We observed that, upon encorafenib withdrawal, both the control and PLEKHA4 knockdown samples started to grow, but to different extents (**Figure 7B**). Further analysis of the tumor xenografts during the 14-day timecourse and at the endpoint confirmed that upon encorafenib withdrawal, both the encorafenib + PLEKHA4 knockdown and encorafenib alone samples had grown, but to different extents (**Figure 7C–D**). Notably, the encorafenib + PLEKHA4 knockdown sample exhibited a slower growth during the relapse phase compared to encorafenib only.

From these data, we conclude that PLEKHA4 knockdown synergizes with BRAF inhibition to prevent melanoma growth in a xenograft model. Further, sustained PLEKHA4 knockdown following encorafenib removal, as a model of resistance/relapse, had a partial but substantial effect on proliferation, suggesting that removal of PLEKHA4 might be therapeutically beneficial in melanoma in combination with existing targeted therapies.

## DISCUSSION

Wnt/β-catenin signaling is a central pathway in embryonic development. In adults, it controls many aspects of cell and tissue homeostasis, including cell proliferation, differentiation, migration (17). Alterations to Wnt signaling that perturb it beyond the normal homeostatic range occur in many diseases; in particular, elevated Wnt signaling occurs in many cancers. In certain instances, mutations to core Wnt components are clearly understood to be drivers of oncogenesis, e.g., in colorectal cancer, where more than 80% of cases feature mutations in adenomatous polyposis coli (APC) that lead to hyperactive Wnt signaling and associated pathogenesis (34). In other cancers with elevated levels of Wnt signaling, the causal nature of this pathway in oncogenesis is not as clear.

Several studies have implicated increased Wnt signaling in melanoma, and yet the functional consequences of this dysregulation in melanoma are not entirely understood (18–20,56). In particular, elevated levels of nuclear β-catenin have been implicated in both increased proliferation but also, unexpectedly, better prognosis, and they are not reliable markers of initial transformation events (20,57). Nuclear β-catenin alone may not necessarily correlate with cellular phenotype, suggesting interplay of additional factors in the regulation of Wnt/β-catenin signaling in melanoma (58). Though the role of Wnt signaling as a sole driver of melanoma progression is controversial, its role in supporting proliferation in certain mutant backgrounds is clearer (21–25). In this context, our study provides important additional evidence implicating Wnt/β-catenin in melanoma proliferation in both BRAF and NRAS mutant backgrounds.

Inhibition of Wnt signaling is a promising route to new anti-cancer therapies, if achievable in a selective or targeted manner that minimizes damage to non-cancerous tissues (19,34–36). Because of challenges associated with targeting core Wnt pathway components, efforts have shifted in recent years toward gaining a deeper understanding of proteins that regulate the strength of Wnt signaling. Among this growing list of modulators, or tuners, PLEKHA4 stands out as a protein with an interesting mechanism of action and potential relevance to melanoma.

Previously, we established that PLEKHA4 enhances Wnt signaling by sequestering and inactivating the Cullin-3 (CUL3) substrate-specific adaptor KLHL12 and prevents DVL polyubiquitination by the CUL3–KLHL12 E3 ubiquitin ligase (40,47). Here, we establish that this fundamental mechanism of tuning Wnt signaling strength could be highly beneficial in the context of melanoma. Melanoma cells express higher levels of PLEKHA4 than more than other 20 cancers investigated by the TCGA (2), and even partial removal of PLEKHA4 by siRNA or shRNA dramatically lowers proliferation and increases apoptosis in vitro and in vivo. PLEKHA4 knockdown exhibits a similar mechanism of action in melanoma cells, i.e., on DVL levels and Wnt signaling strength, as well as strong effects on clonogenic capacity in vitro. In tumor xenograft and allograft models using both BRAF- and NRAS-mutant melanomas, removal of PLEKHA4 by shRNA prevented tumor growth. And finally, in a BRAF-mutant melanoma, PLEKHA4 shRNA synergized with a clinically used BRAF inhibitor, leading to much stronger tumor shrinkage, and its positive effects help to keep growth slow even after removal of the inhibitor, as a model of resistance.

These results showing synergy with a BRAF inhibitor reinforce that, while MAP kinase signaling is a predominant player in melanoma, Wnt/β-catenin plays important roles in supporting proliferation. Other modulators of Wnt signaling affect melanoma proliferation. For example, Dkk-1, a negative regulator of Wnt signaling, exhibits reduced expression in melanoma, and its activation inhibits tumorigenicity and induces apoptosis in melanoma (59,60). Another negative regulator of Wnt signaling, WIF-1 (Wnt inhibitory factor-1), is downregulated in melanoma progression (61). Both MAP kinase and Wnt/β-catenin signaling regulate the activity of MITF, a master regulator of melanoma progression in both BRAF and NRAS mutant backgrounds (62–64).

Our results underscore Wnt/β-catenin signaling, and its regulator PLEKHA4, as important players controlling proliferation in both BRAF- and NRAS-mutant melanomas. PLEKHA4 knockdown in melanoma cells strongly affected levels of the canonical Wnt/β-catenin targets Cyclin D1 and c-Myc, which ensure progression through the G1/S cell cycle transition. Disruption of Wnt signaling via other means (DVL knockdown or global pharmacological inhibition of Wnt production) resulted in similar phenotypes to PLEKHA4 knockdown. Thus, PLEKHA4 is critical for ensuring adequate canonical Wnt/β-catenin signaling to maintain a proliferative phenotype in melanoma.

Further, these results suggest that PLEKHA4 inhibition might be therapeutically beneficial in both NRAS-mutant melanomas, for which there are no targeted therapies, and for BRAF-mutant melanomas, where PLEKHA4 inhibition could be investigated in combination with existing BRAF or MEK inhibitors. In principle, PLEKHA4 inhibition might also synergize with immunotherapies, which would represent an interesting future direction.

PLEKHA4 is not a canonical drug target. It is a multidomain adaptor protein, not a receptor, ion channel, or enzyme. Yet, our previous work sheds light on several protein-lipid and protein-protein interactions that could be targeted (47). Its tripartite N-terminal region, which includes a pleckstrin homology (PH) domain, binds to anionic phosphoinositides to localize the protein to the plasma membrane. C-terminal coiled-coil and intrinsically disordered regions mediate oligomerization into membrane-associated clusters that are potentially phase-separated. A central proline-rich domain binds to KLHL12, and all three of these molecular elements (lipid binding, oligomerization, and KLHL12 binding) are featured in its mechanism of action to prevent DVL ubiquitination and enhance Wnt signaling.

In principle, small-molecule ligands could be developed to target the phosphoinositide binding site of the PH domain (65–67) or disrupt interactions between the proline-rich domain and KLHL12 (68) or homotypic interactions involved in oligomerization and cluster formation (69). Further, ligands that bind to PLEKHA4 but do not disrupt function could still serve as starting points for development of PROTACs/degraders (70,71). Finally, a global knockout of the *Drosophila* ortholog of PLEKHA4, *kramer*, is viable (47), raising the possibility that mammalian PLEKHA4 may be dispensable for development and less critical for maintaining homeostatic Wnt signaling. Yet, this study implicates it as a vulnerability for melanoma cells. Thus, we believe that PLEKHA4 defines a new type of drug target for melanoma.

Interestingly, our previous work on PLEKHA4 and *kramer* established that these proteins can also mediate non-canonical, β-catenin-independent Wnt signaling (47). In particular, in *Drosophila*, *kramer* knockout resulted in defects in planar cell polarity through effects on *dishevelled*, a pathway that shares key aspects with mammalian non-canonical Wnt signaling, including profound effects on the actin cytoskeleton (72). In melanoma, non-canonical Wnt signaling is implicated in a migratory phenotype, whereas canonical Wnt/β-catenin signaling controls proliferation. Melanoma progression has been described to involve a phenotype switching scenario, wherein alternating cycles of proliferation and migration lead to disease spread and eventually to metastasis (58).

Crucially, DVL is a central signaling molecule in both the canonical and non-canonical pathways (39), and thus it is not surprising that PLEKHA4, which regulates DVL levels, has the potential to affect multiple types of Wnt signaling, depending on the context (47). In the in vitro and xenograft models here, which are geared toward evaluation of the proliferative stages of melanoma, we found a strong effect on removal of PLEKHA4. Examination of effects of PLEKHA4 removal on non-canonical Wnt signaling in the context of a migratory phenotype represents an interesting future direction and could reveal that a single protein, PLEKHA4, might be relevant in suppressing later stages of melanoma, including metastasis, where the cancer cells exhibit an invasive phenotype.

In summary, we have identified PLEKHA4 as an important mediator of a proliferative phenotype in BRAF- and NRAS-mutant melanoma. We demonstrate that PLEKHA4 acts in this role as a positive regulator of Wnt/β-catenin signaling in this context, helping to clarify the role of Wnt/β-catenin signaling in this context and revealing another layer of regulation in the Wnt/β-catenin signaling axis that controls the G1/S cell cycle transition to maintain melanoma proliferation.

## MATERIALS AND METHODS

### Cell culture

WM266-4 (NCI PSOC) and SK-MEL-2 (NCI PSOC) cells were cultured in minimum essential medium (MEM, Corning), L Wnt-3a cells and control L cells (ATCC) were cultured in Dulbecco’s modified Eagle medium (DMEM, Corning) and YUMM1.7 (ATCC) cells were cultured in Dulbecco’s modified Eagle medium/ Ham’s F-12 medium (DMEM/F-12, Corning) supplemented with 10% fetal bovine serum (FBS, Corning) and 1% penicillin/streptomycin (P/S, Corning) at 37 °C in a 5% CO2 atmosphere. Stable expression of doxycycline-inducible shRenilla control, shPLEKHA4 hairpins or mouse shPLEKHA4 hairpins in WM266-4 or SK-MEL-2 (human) and YUMM1.7 (mouse) cells was achieved by transducing above-mentioned hairpin plasmids cloned in LT3GEPIR vector (a gift from Lewis Cantley) as previously described (73). Stable expression of cell cycle indicator plasmid pLenti6.2-FUCCI (Fluorescence Ubiquitination Cell Cycle Indicator, a gift from Jan Lammerding) was achieved by transducing FUCCI plasmid into WM266-4 and SK-MEL-2 cells. After transduction (48 h), hairpin transduced cells were selected with 2.5 μg/mL puromycin (Sigma-Aldrich) and FUCCI plasmid-transduced cells were selected with 2 μg/mL blasticidin (Alfa Aesar). Upon completion of drug selection, FUCCI transduced WM266-4 and SK-MEL-2 cells were sorted using FACS to ensure 99.9% of fluorescent cell population before use. Conditioned media (CM) from L and L Wnt-3a cells was harvested as previously described (47). Cell lines were obtained and used without further authentication.

### Animal husbandry

All mice used for experiments were approved by Center for Animal Resources and Education (CARE) facility at Cornell University. The C57BL/6J mice were purchased from Jackson laboratory and NSG (NOD *scid* gamma) mice were purchased from the PATh PDX facility at Cornell University. The animals were housed and bred on a 12 h light and dark cycle. C57BL/6J mice were used for xenografting/allografting the YUMM1.7 syngeneic mouse cell line, and NSG mice were used for xenografting WM266-4 or SK-MEL-2 (human melanoma) cell lines. Mice were euthanized when tumors reached the maximum size allowed by the approved animal protocol accounting for the animal’s health and mobility.

### Plasmids and cloning

The cloning for GFP and siRNA-resistant PLEKHA4-GFP plasmids for rescue experiments have been described previously (47). Viral transduction vector pCDH-mCherry-Blasticidin was obtained as a gift from Jan Lammerding’s lab at Cornell University. The vector was digested using standard cloning procedure with EcoRI and NotI to remove mCherry. The full length GFP or siRNA resistant PLEKHA4-GFP PCR fragments were subcloned into the digested pCDH vector. For cloning pCDH-DVL2-GFP and pCDH-DVL3-GFP, full length DVL2 and DVL3 PCR-fragments were amplified from 3X-FLAG-DVL2 (Addgene #24802) and XE251-pcDNA3.1 (zeo) FLAG-hDsh3 (Addgene #16758) respectively. pCDH-PLEKHA4-GFP was digested with EcoRI and AgeI to remove PLEKHA4 fragment and the remaining vector was used to subclone the DVL2 or DVL3 fragments.

For generation of doxycycline-inducible stable shPLEKHA4 lines, 12 each human or mouse shRNA constructs against PLEKHA4 was cloned into LT3GEPIR vector as described previously (73). All constructs were verified by Sanger sequencing (Cornell University Biotechnology Resource Center Genomics Facility).

### Transfection of siRNAs

DsiRNA duplexes were obtained from Integrated DNA Technologies. Transfections with siRNA were performed using Lipofectamine RNAiMAX with the appropriate duplexes (see **Table 1**) as described previously (47), and 48 h post transfection, cells were analyzed via Western blot, flow cytometry or other readouts.

### Virus production

Exogenous protein expression was achieved in WM266-4 and SK-MEL-2 cells via lentiviral transduction. Lentivirus was produced in HEK 293TN cells, which were seeded to achieve a 90% confluency on the day of transfection with the viral plasmids. Packaging plasmids VSVg and PAX2, along with the lentiviral plasmid, were transfected overnight in the ratio of 1:3.1:4.2 in the HEK 293TN cells using Lipofectamine 2000. Fresh media was changed the next morning and the transfection media was discarded. Thirty-six h post transfection, the lentivirus-containing media was collected, and cells were replenished with new fresh media. The media collection was performed every 8 h for a total of four times. The lentivirus-containing media was filtered through 0.45 μm syringe filters and stored in 4 °C until use. The lentivirus media was used within two weeks after production. For production of shRNA-containing lentivirus, the same protocol was used, except that packaging plasmids VSVg and PAX2, along with shRNA plasmids, were mixed and transfected overnight in the ratio of 1:1.8:3.7.

### Lentivirus transduction of plasmids

Depending on the experiment, cells were seeded 1–2 d prior to viral transduction. On the day of transduction, media was aspirated, and one part of fresh media was added with 8 μg/mL of Polybrene (Millipore) and spread evenly. Three parts of filtered lentivirus media was added and gently mixed. For the highest transduction efficiency, the process was repeated four times every 10-12 h. For stable cell line generation, cells were selected with the appropriate selection drug as described above. For all other experiments, cells were used without selection. Transduction efficiency was determined using fluorescence microscopy for every lentivirus transduction experiment using Hoechst 33342 (Thermo Fisher) as the counterstain. The transduction efficiency for these experiments were determined to be 80–90%. Images were acquired on a Zeiss LSM 800 confocal laser scanning microscope equipped with Plan Apochromat objectives (20x 0.8 NA or 40x 1.4 NA), and two GaAsP PMT detectors. Solid-state lasers (405, 488, 561, and 640 nm) were used to excite blue, green, red and far-red fluorescence respectively. Images were acquired using Zeiss Zen Blue 2.3 (confocal) and analyzed using ImageJ.

### Cell proliferation assay

SiRNA duplexes (50 nM) against PLEKHA4, DVL2 and DVL3 was performed overnight on either WM266-4 or SK-MEL-2 cells on a 6-well plate. Sixteen h post RNAi, cells were lifted using trypsin and counted three times using hematocytometer. Four thousand cells were seeded in each well in a low evaporation lid 96-well plate. To assess the effect of Wnt inhibition on proliferation of WM266-4 or SK-MEL-2, cells were counted and seeded in the 96-well plate with media containing either DMSO vehicle or 2.5 μM IWP-4 (Inhibitor of Wnt Production-4). Images were acquired every 1 h for at least 4 d using 20X objective in an IncuCyte cell incubator.

### Anchorage-dependent colony formation assay

SiRNA duplexes (50 nM) against PLEKHA4, DVL2 and DVL3 was performed overnight on either WM266-4 or SK-MEL-2 cells on a 6-well plate. Sixteen h post RNAi, cells were lifted using trypsin, counted three times using hematocytometer and 4000 cells were plated and dispersed evenly in each well of a 6-well plate. Fresh media was changed every 3 d. For Wnt inhibition experiments, 4000 untreated cells were plated in a 6-well in either DMSO control or IWP-4 containing media. The cells were grown for two weeks and stopped once colonies were observed. At the end of two weeks, cells were washed with PBS, fixed with methanol for 1 h at room temperature and stained overnight with 0.1% crystal violet in 95% ethanol. The lids were propped slightly open to let the stain solution completely evaporate. The next day, plates were rinsed gently with cold water to wash off the excess stain and dried with lids open for approximately 3 h. Images were acquired with a Bio-Rad gel scanning doc and colonies were counted using ImageJ.

### Anchorage-independent soft agar assay

SiRNA duplexes (50 nM) against PLEKHA4, DVL2 and DVL3 was performed overnight on either WM266-4 or SK-MEL-2 cells on a 6-well plate. Sixteen h post RNAi, cells were lifted using trypsin, counted three times using hematocytometer and 5000 cells were plated and dispersed evenly in each well of a 6-well plate. Soft agar assay was set up following the protocol as described previously (54). Three weeks after the seeding, colonies were observed and stained overnight in a 37 °C incubator with 1 mg/mL nitrotetrazolium blue solution in 1X PBS. Images were acquired with a Bio-Rad gel scanning doc and colonies were counted using ImageJ.

### Cell cycle analysis

#### Unsynchronized

To initially assess the molecular defects in proliferation, cell cycle analysis was performed in unsynchronized WM266-4 cells. SiRNA duplexes (50 nM) against PLEKHA4 was performed overnight on a 12-well plate. Forty-eight h post RNAi, cells were lifted using trypsin, fixed overnight with prechilled ethanol and stained using propidium iodide following the protocol as described previously (74). Cells were analyzed via flow cytometry.

#### Synchronized

The unsynchronized WM266-4 cells showed a defect in G1/S cell cycle transition. To quantify this defect, stable cells containing the genetically encoded cell cycle indicator plasmid FUCCI were generated as described above. Cells were seeded on a 15-cm dish and grown until confluency was 90%. They were starved using media without FBS for 48 h and siRNA duplexes (50 nM) against PLEKHA4 was performed overnight in the starvation media at the end of the 48-h period. Sixteen h post RNAi, cells were stimulated using fresh media with FBS for 36 h. Cells were lifted using trypsin and fixed overnight with pre-chilled ethanol at 4 °C. The next day, cells were washed three times with FACS buffer (0.1% FBS in 1X PBS), analyzed via flow cytometry and quantified for G1 (red fluorescent) vs. S-G2-M (green fluorescent) cell populations. Quantifications are from at least three biological replicates.

#### Rescue

Virus-containing media containing each rescue construct (GFP, PLEKHA4-GFP, DVL2-GFP, or DVL3-GFP) was generated as described above and used to rescue the G1/S transition defect. RNAi against PLEKHA4 was performed as described on a 60-mm plate, and 16 h post RNAi, cells were stimulated with the rescue media (a mix of 1.5 mL fresh media and 2.5 mL virus-containing rescue media) following the virus transduction protocol. Forty-eight h post RNAi, cells were harvested and fixed overnight with prechilled ethanol at 4 °C. The rescue was performed orthogonally using both the propidium iodide cell cycle analysis method on wild-type, non-fluorescent cells and the FUCCI stable fluorescent cell lines. Cells were analyzed via flow cytometry and quantified for G1 cell populations.

### Tumor xenografts

NSG and C57BL/6J mice were bred as described above. Stable cell lines with doxycycline-inducible hairpins against PLEKHA4 or control were generated as described above. A day before the tumor xenograft injections, the dorsal side of 4-6 weeks old mice was shaved to enable four injections on each animal, two each near the upper and lower flanks. On the day of xenograft, cells were lifted using trypsin, counted three times using a hematocytometer, and resuspended in media containing 1% penicillin/streptomycin. A 1:1 mixture of cells:Matrigel was made, and 1×10^6^ of shRNA-expressing WM266-4 or SK-MEL-2 cells were subcutaneously injected into NSG mice using a 28-gauge needle. The same procedure was used for shRNA-expressing YUMM1.7 cells except that 1×10^5^ cells were injected subcutaneously into C57BL/6J mice. All injections were performed within 30 min of preparing the cells/Matrigel mix. After injection, the mice were monitored every 2 d. For WM266-4 and YUMM1.7 xenografts, tumor formation appeared around day 12, whereas for SK-MEL-2, the tumor formation appeared after 1.5 months post injection. Doxycycline (1 mg/mL in sterile water) was administered in the drinking water in amber bottles and changed every 2 d for 12 d total for WM266-4, 10 d total for YUMM1.7, and for 16 d total for SK-MEL-2 xenografts. Tumor progression was monitored every 2 d by measuring the tumor dimensions using a digital caliper and calculating the volume using the equation, v = 0.5233*l*w^2^. The volumes were plotted, and statistical analysis was performed.

### Tumor xenografts to assess synergy between BRAFi and shPLEKHA4

NSG mice and cells were treated as described above. On the day of xenograft, 1×10^6^ of shRNA-expressing WM266-4 or control cells were injected subcutaneously. After injection, the mice were monitored every 2 d and tumor formation was appeared around day 12. On day 12, the synergy treatment was started by randomly selecting the mice for control vs. BRAF inhibition along with shRNA induction. Doxycycline (1 mg/mL in sterile water) was administered every 2 d to induce the shRNA, and encorafenib (BRAF inhibitor, 30 mg/kg in 0.5% carboxymethylcellulose and 0.05% Tween-80 in 1X PBS) was given via oral gavage every day, for 12 d. The solution was made fresh every day. For control animals, doxycycline and DMSO in vehicle solution was administered. Tumor progression was monitored as described above. Tumor relapse upon encorafenib withdrawal in presence and absence of PLEKHA4 knockdown was assessed. At the end of 12 d of encorafenib treatment, the drug was withdrawn but doxycycline was continued for another 14 d. Tumor progression was monitored every 2 days, and volumes were plotted for statistical analysis.

### Western blot analysis of DVL, Myc, and Cyclin D1 levels

SiRNA duplexes (50 nM) against PLEKHA4, DVL2, or DVL3 were used to perform knockdown on either WM266-4 or SK-MEL-2 cells on a 6-well plate. Forty-eight h post-RNAi, cells were harvested and analyzed by Western blot as described previously (47). The levels of DVL2, DVL3, Myc, and Cyclin D1 were quantified. Reported quantifications are from at least three biological replicates. For analysis of endogenous Axin2 levels, siRNA duplexes (50 nM) against PLEKHA4 were transfected into WM266-4 or SK-MEL-2 cells on a 6-well plate. Cells were stimulated with Wnt3a-conditioned media, harvested, analyzed by Western blot, and and Axin2 levels were quantified as described previously (47). For rescue experiments, samples were generated as described above for cell cycle rescue analysis. Forty-eight h post-RNAi, cells were harvested, normalized and analyzed via Western blot for DVL2, DVL3, Myc, and Cyclin D1 levels. Reported quantifications are from at least three biological replicates.

### Luciferase Wnt reporter assay

#### Generation of Wnt reporter WM266-4 and SK-MEL-2 cells stably expressing Firefly and Renilla luciferase

WM266-4 or SK-MEL-2 cells were co-transduced with conditioned media containing lentiviruses bearing Firefly luciferase-7TFP (Addgene #24308, β-catenin reporter) and Renilla luciferase pLenti.PGK.blast-Renilla Luciferase (Addgene #74444) as described above. After 48 h, cells were treated with 2.5 μg/mL puromycin dihydrochloride (Sigma-Aldrich) and 2 μg/mL blasticidin S hydrochloride (Alfa Aesar) until the appearance of resistant colonies. The Wnt/β-catenin-dependent luciferase reporter WM266-4 and SK-MEL-2 cell lines were tested for luciferase activity and was used for the siRNA-based Wnt luciferase reporter assay.

#### Transient knockdown assay

SiRNA-mediated knockdown was performed against PLEKHA4 in Wnt/β-catenin dependent reporter WM266-4 and SK-MEL-2 cells on 6-well plates. After 30 h of cell growth post-transfection, cells were treated with sterile-filtered Wnt3a-containing conditioned media in a 1:1 ratio with fresh media for 30 h. Cells were then lysed, and 150 μL of lysates were transferred to an opaque 96-well flat-bottom plate (Greiner) for measuring chemiluminescence. 50 μL of luciferin (firefly luciferase substrate) was added to each well, and the firefly luciferase signal was read by a Tecan plate reader. Subsequently, 50 μL of Renilla luciferase substrate, which contains 25 μM of firefly luciferase inhibitor 4-(6-methyl-1,3-benzothiazol-2-yl)-aniline (Enamine.net), was added to each well, and the Renilla luciferase signal was read.

#### Stable Knockdown

ShRNAs against PLEKHA4 in WM266-4 and SK-MEL-2 cells were induced by addition of 2.5 μg/mL doxycycline for 10 d in 6-well plates. Doxycycline-containing media was exchanged for fresh media every 2 d. On day 8, the cells were treated with a 1:1:1 mixture of 7TFP lentivirus-containing conditioned media:PGK-Renilla lentivirus-containing conditioned media:fresh media, along with 8 μg/mL polybrene and 2.5 μg/mL doxycycline for 24 h. Spent media was exchanged for fresh 1:1:1 media mixture as described above every 12 h. On day 9, the cells were induced by adding Wnt3a-containing conditioned media in a 1:1 ratio with fresh media containing doxycycline for 30 h. Firefly and Renilla luciferase signals were then obtained as described above.

### Statistics and reproducibility

All experiments were performed in at least three biological replicates. Imaging figures show representative images from each experiment. For experiments involving quantification of comparisons between two groups, statistical significance was calculated using unpaired two-tailed Student’s t-test with unequal variance in Excel or GraphPad Prism. For experiments involving quantification of comparisons between more than two experimental groups, statistical significance was calculated using a one-way ANOVA with post-hoc Tukey test in R. The number of biological replicates analyzed is stated in the legend, and statistical significance of p < 0.05 or lower is reported. All raw data were plotted using either Excel or GraphPad Prism. In figures containing bar graphs, the height of the bar is the mean, the error bars represent standard deviation, and each overlaid dot represents an individual biological replicate. In figures containing scatter plots, the black line is the mean, and each dot represents an individual biological replicate. In figures containing IncuCyte proliferation data and tumor xenograft progression data, the means at various time points were plotted, and error bars represent standard deviation. Tumor xenograft progression measurements were performed in a blinded manner. In Figure 1A, the boxes represent the middle quartiles, with the line in the middle denoting the median.

## Data availability statement

The authors declare that all data supporting the findings of this study are available within the paper and its supporting information files.

## ACKNOWLEDGMENTS

This work was supported by the NIH (R01GM131101) and the Alfred P. Sloan Foundation (Sloan Research Fellowship to J.M.B.). X.C. was supported by a Cornell Fellowship. We thank Dahihm Kim, Tyler Kirby, Hyeongsun Moon, and Diana Wang for technical help, and the Cantley, Emr, Fromme, Lammerding, and Yu labs for sharing reagents and equipment.

## Notes

### Competing Interest Statement

The authors have declared no competing interest.

## REFERENCES

1. Miller AJ, Mihm MC. Melanoma. N Engl J Med [Internet]. Massachusetts Medical Society; 2006;355:51–65. Available from: https://doi.org/10.1056/NEJMra052166

2. Akbani R, Akdemir KC, Aksoy BA, Albert M, Ally A, Amin SB, et al. Genomic Classification of Cutaneous Melanoma. Cell. 2015;161:1681–96.

3. Hayward NK, Wilmott JS, Waddell N, Johansson PA, Field MA, Nones K, et al. Whole-genome landscapes of major melanoma subtypes. Nature [Internet]. 2017;545:175–80. Available from: https://doi.org/10.1038/nature22071

4. Flaherty KT, Puzanov I, Kim KB, Ribas A, McArthur GA, Sosman JA, et al. Inhibition of Mutated, Activated BRAF in Metastatic Melanoma. N Engl J Med [Internet]. Massachusetts Medical Society; 2010;363:809–19. Available from: https://doi.org/10.1056/NEJMoa1002011

5. Poulikakos PI, Persaud Y, Janakiraman M, Kong X, Ng C, Moriceau G, et al. RAF inhibitor resistance is mediated by dimerization of aberrantly spliced BRAF(V600E). Nature [Internet]. 2011;480:387–90. Available from: https://doi.org/10.1038/nature10662

6. Bollag G, Hirth P, Tsai J, Zhang J, Ibrahim PN, Cho H, et al. Clinical efficacy of a RAF inhibitor needs broad target blockade in BRAF-mutant melanoma. Nature [Internet]. 2010;467:596–9. Available from: https://doi.org/10.1038/nature09454

7. Long G V, Stroyakovskiy D, Gogas H, Levchenko E, de Braud F, Larkin J, et al. Combined BRAF and MEK Inhibition versus BRAF Inhibition Alone in Melanoma. N Engl J Med [Internet]. Massachusetts Medical Society; 2014;371:1877–88. Available from: https://doi.org/10.1056/NEJMoa1406037

8. Larkin J, Ascierto PA, Dréno B, Atkinson V, Liszkay G, Maio M, et al. Combined Vemurafenib and Cobimetinib in BRAF-Mutated Melanoma. N Engl J Med [Internet]. Massachusetts Medical Society; 2014;371:1867–76. Available from: https://doi.org/10.1056/NEJMoa1408868

9. Nazarian R, Shi H, Wang Q, Kong X, Koya RC, Lee H, et al. Melanomas acquire resistance to B-RAF(V600E) inhibition by RTK or N-RAS upregulation. Nature [Internet]. 2010;468:973–7. Available from: https://doi.org/10.1038/nature09626

10. Rizos H, Menzies AM, Pupo GM, Carlino MS, Fung C, Hyman J, et al. BRAF Inhibitor Resistance Mechanisms in Metastatic Melanoma: Spectrum and Clinical Impact. Clin Cancer Res [Internet]. 2014;20:1965 LP – 1977. Available from: http://clincancerres.aacrjournals.org/content/20/7/1965.abstract

11. Lito P, Rosen N, Solit DB. Tumor adaptation and resistance to RAF inhibitors. Nat Med [Internet]. 2013;19:1401–9. Available from: https://doi.org/10.1038/nm.3392

12. Hodi FS, O’Day SJ, McDermott DF, Weber RW, Sosman JA, Haanen JB, et al. Improved Survival with Ipilimumab in Patients with Metastatic Melanoma. N Engl J Med [Internet]. Massachusetts Medical Society; 2010;363:711–23. Available from: https://doi.org/10.1056/NEJMoa1003466

13. Hamid O, Robert C, Daud A, Hodi FS, Hwu W-J, Kefford R, et al. Safety and Tumor Responses with Lambrolizumab (Anti–PD-1) in Melanoma. N Engl J Med [Internet]. Massachusetts Medical Society; 2013;369:134–44. Available from: https://doi.org/10.1056/NEJMoa1305133

14. Hu-Lieskovan S, Mok S, Homet Moreno B, Tsoi J, Robert L, Goedert L, et al. Improved antitumor activity of immunotherapy with BRAF and MEK inhibitors in *BRAF*^V600E^ melanoma. Sci Transl Med [Internet]. 2015;7:279ra41 LP–279ra41. Available from: http://stm.sciencemag.org/content/7/279/279ra41.abstract

15. Zhan T, Rindtorff N, Boutros M. Wnt signaling in cancer. Oncogene [Internet]. Nature Publishing Group; 2017;36:1461–73. Available from: http://dx.doi.org/10.1038/onc.2016.304

16. Clevers H, Nusse R. Wnt/β-catenin signaling and disease. Cell [Internet]. Cell Press; 2012 [cited 2018 Feb 1];149:1192–205. Available from: https://www.sciencedirect.com/science/article/pii/S0092867412005867

17. MacDonald BT, Tamai K, He X. Wnt/β-Catenin Signaling: Components, Mechanisms, and Diseases. Dev Cell [Internet]. Elsevier Inc.; 2009;17:9–26. Available from: http://dx.doi.org/10.1016/j.devcel.2009.06.016

18. Webster MR, Weeraratna AT. A Wnt-er Migration: The Confusing Role of β-Catenin in Melanoma Metastasis. Sci Signal [Internet]. 2013;6:pe11 LP-pe11. Available from: http://stke.sciencemag.org/content/6/268/pe11.abstract

19. Jackstadt R, Hodder MC, Sansom OJ. WNT and β-Catenin in Cancer: Genes and Therapy. Annu Rev Cancer Biol [Internet]. Annual Reviews; 2020;4:177–96. Available from: https://doi.org/10.1146/annurev-cancerbio-030419-033628

20. Gajos-Michniewicz A, Czyz M. Wnt Signaling in Melanoma. Int J Mol Sci. 2020;21:4852.

21. Delmas V, Beermann F, Martinozzi S, Carreira S, Ackermann J, Kumasaka M, et al. β-Catenin induces immortalization of melanocytes by suppressing p16INK4a expression and cooperates with N-Ras in melanoma development. Genes Dev [Internet]. 2007;21:2923–35. Available from: http://genesdev.cshlp.org/content/21/22/2923.abstract

22. Pawlikowski JS, McBryan T, van Tuyn J, Drotar ME, Hewitt RN, Maier AB, et al. Wnt signaling potentiates nevogenesis. Proc Natl Acad Sci USA [Internet]. 2013;110:16009 LP – 16014. Available from: http://www.pnas.org/content/110/40/16009.abstract

23. Juan J, Muraguchi T, Iezza G, Sears RC, McMahon M. Diminished WNT -> β-catenin -> c-MYC signaling is a barrier for malignant progression of BRAFV600E-induced lung tumors. Genes Dev [Internet]. 2014/03/03. Cold Spring Harbor Laboratory Press; 2014;28:561–75. Available from: https://pubmed.ncbi.nlm.nih.gov/24589553

24. Damsky WE, Curley DP, Santhanakrishnan M, Rosenbaum LE, Platt JT, Gould Rothberg BE, et al. β-Catenin Signaling Controls Metastasis in Braf-Activated Pten-Deficient Melanomas. Cancer Cell [Internet]. 2011;20:741–54. Available from: http://www.sciencedirect.com/science/article/pii/S1535610811004053

25. Sun Q, Lee W, Mohri Y, Takeo M, Lim CH, Xu X, et al. A novel mouse model demonstrates that oncogenic melanocyte stem cells engender melanoma resembling human disease. Nat Commun [Internet]. Springer US; 2019;10:1–16. Available from: http://dx.doi.org/10.1038/s41467-019-12733-1

26. Chien AJ, Haydu LE, Biechele TL, Kulikauskas RM, Rizos H, Kefford RF, et al. Targeted BRAF Inhibition Impacts Survival in Melanoma Patients with High Levels of Wnt/β-Catenin Signaling. PLoS One [Internet]. Public Library of Science; 2014;9:e94748. Available from: https://doi.org/10.1371/journal.pone.0094748

27. Kageshita T, Hamby C V, Ishihara T, Matsumoto K, Saida T, Ono T. Loss of β-catenin expression associated with disease progression in malignant melanoma. Br J Dermatol [Internet]. John Wiley & Sons, Ltd; 2001;145:210–6. Available from: https://doi.org/10.1046/j.1365-2133.2001.04336.x

28. Bachmann IM, Straume O, Puntervoll HE, Kalvenes MB, Akslen LA. Importance of P-Cadherin, β-Catenin, and Wnt5a/Frizzled for Progression of Melanocytic Tumors and Prognosis in Cutaneous Melanoma. Clin Cancer Res [Internet]. 2005;11:8606 LP – 8614. Available from: http://clincancerres.aacrjournals.org/content/11/24/8606.abstract

29. Chien AJ, Moore EC, Lonsdorf AS, Kulikauskas RM, Rothberg BG, Berger AJ, et al. Activated Wnt/ß-catenin signaling in melanoma is associated with decreased proliferation in patient tumors and a murine melanoma model. Proc Natl Acad Sci USA [Internet]. 2009;106:1193 LP – 1198. Available from: http://www.pnas.org/content/106/4/1193.abstract

30. Xue G, Romano E, Massi D, Mandalà M. Wnt/β-catenin signaling in melanoma: Preclinical rationale and novel therapeutic insights. Cancer Treat Rev. 2016;49:1–12.

31. Webster MR, Xu M, Kinzler KA, Kaur A, Appleton J, O’Connell MP, et al. Wnt5A promotes an adaptive, senescent-like stress response, while continuing to drive invasion in melanoma cells. Pigment Cell Melanoma Res [Internet]. John Wiley & Sons, Ltd; 2015;28:184–95. Available from: https://doi.org/10.1111/pcmr.12330

32. Anastas JN, Kulikauskas RM, Tamir T, Rizos H, Long G V, von Euw EM, et al. WNT5A enhances resistance of melanoma cells to targeted BRAF inhibitors. J Clin Invest [Internet]. The American Society for Clinical Investigation; 2014;124:2877–90. Available from: https://doi.org/10.1172/JCI70156

33. Hoek KS, Eichhoff OM, Schlegel NC, Döbbeling U, Kobert N, Schaerer L, et al. In vivo Switching of Human Melanoma Cells between Proliferative and Invasive States. Cancer Res [Internet]. 2008;68:650 LP – 656. Available from: http://cancerres.aacrjournals.org/content/68/3/650.abstract

34. Nusse R, Clevers H. Wnt/β-Catenin Signaling, Disease, and Emerging Therapeutic Modalities. Cell [Internet]. Cell Press; 2017 [cited 2018 Feb 1];169:985–99. Available from: https://www.sciencedirect.com/science/article/pii/S0092867417305470

35. Krishnamurthy N, Kurzrock R. Targeting the Wnt/beta-catenin pathway in cancer: Update on effectors and inhibitors. Cancer Treat Rev. 2018;62:50–60.

36. Tran FH, Zheng JJ. Modulating the wnt signaling pathway with small molecules. Protein Sci [Internet]. John Wiley & Sons, Ltd; 2017;26:650–61. Available from: https://doi.org/10.1002/pro.3122

37. Kahn M. Can we safely target the WNT pathway? Nat Rev Drug Discov [Internet]. 2014;13:513–32. Available from: https://doi.org/10.1038/nrd4233

38. Mlodzik M. The Dishevelled Protein Family: Still Rather a Mystery After Over 20 Years of Molecular Studies [Internet]. 1st ed. Curr. Top. Dev. Biol. Elsevier Inc.; 2016. Available from: http://dx.doi.org/10.1016/bs.ctdb.2015.11.027

39. Gao C, Chen YG. Dishevelled: The hub of Wnt signaling. Cell Signal [Internet]. Elsevier Inc.; 2010;22:717–27. Available from: http://dx.doi.org/10.1016/j.cellsig.2009.11.021

40. Angers S, Thorpe CJ, Biechele TL, Goldenberg SJ, Zheng N, MacCoss MJ, et al. The KLHL12–Cullin-3 ubiquitin ligase negatively regulates the Wnt–β-catenin pathway by targeting Dishevelled for degradation. Nat Cell Biol [Internet]. 2006;8:348–57. Available from: http://www.nature.com/doifinder/10.1038/ncb1381

41. Nielsen CP, Jernigan KK, Diggins NL, Webb DJ, MacGurn JA. USP9X Deubiquitylates DVL2 to Regulate WNT Pathway Specification. Cell Rep [Internet]. Elsevier; 2019;28:1074–1089.e5. Available from: https://doi.org/10.1016/j.celrep.2019.06.083

42. Schwarz-Romond T, Fiedler M, Shibata N, Butler PJG, Kikuchi A, Higuchi Y, et al. The DIX domain of Dishevelled confers Wnt signaling by dynamic polymerization. Nat Struct Mol Biol [Internet]. 2007;14:484–92. Available from: https://doi.org/10.1038/nsmb1247

43. Tauriello DVF, Haegebarth A, Kuper I, Edelmann MJ, Henraat M, Canninga-van Dijk MR, et al. Loss of the Tumor Suppressor CYLD Enhances Wnt/β-Catenin Signaling through K63-Linked Ubiquitination of Dvl. Mol Cell [Internet]. 2010;37:607–19. Available from: http://www.sciencedirect.com/science/article/pii/S1097276510001243

44. Ding Y, Zhang Y, Xu C, Tao QH, Chen YG. HECT domain-containing E3 ubiquitin ligase NEDD4L negatively regulates Wnt signaling by targeting dishevelled for proteasomal degradation. J Biol Chem. 2013;288:8289–98.

45. Gao C, Cao W, Bao L, Zuo W, Xie G, Cai T, et al. Autophagy negatively regulates Wnt signalling by promoting Dishevelled degradation. Nat Cell Biol. Nature Publishing Group; 2010;12:781–90.

46. Wei W, Li M, Wang J, Nie F, Li L. The E3 Ubiquitin Ligase ITCH Negatively Regulates Canonical Wnt Signaling by Targeting Dishevelled Protein. Mol Cell Biol [Internet]. 2012;32:3903–12. Available from: http://mcb.asm.org/cgi/doi/10.1128/MCB.00251-12

47. Shami Shah A, Batrouni AG, Kim D, Punyala A, Cao W, Han C, et al. PLEKHA4/kramer Attenuates Dishevelled Ubiquitination to Modulate Wnt and Planar Cell Polarity Signaling. Cell Rep [Internet]. ElsevierCompany.; 2019;27:2157–2170.e8. Available from: https://doi.org/10.1016/j.celrep.2019.04.060

48. Hruz T, Laule O, Szabo G, Wessendorp F, Bleuler S, Oertle L, et al. Genevestigator V3: A Reference Expression Database for the Meta-Analysis of Transcriptomes. Adv Bioinformatics. 2008;2008:1–5.

49. Koelblinger P, Thuerigen O, Dummer R. Development of encorafenib for BRAF-mutated advanced melanoma. Curr Opin Oncol. 2018;30:125–33.

50. Chen B, Dodge ME, Tang W, Lu J, Ma Z, Fan C-W, et al. Small molecule-mediated disruption of Wnt-dependent signaling in tissue regeneration and cancer. Nat Chem Biol. 2009;5:100–7.

51. Sakaue-Sawano A, Kurokawa H, Morimura T, Hanyu A, Hama H, Osawa H, et al. Visualizing Spatiotemporal Dynamics of Multicellular Cell-Cycle Progression. Cell. 2008;132:487–98.

52. Shtutman M, Zhurinsky J, Simcha I, Albanese C, D’Amico M, Pestell R, et al. The cyclin D1 gene is a target of the β-catenin/LEF-1 pathway. Proc Natl Acad Sci USA [Internet]. 1999;96:5522 LP – 5527. Available from: http://www.pnas.org/content/96/10/5522.abstract

53. He T-C, Sparks AB, Rago C, Hermeking H, Zawel L, da Costa LT, et al. Identification of c-MYC as a Target of the APC Pathway. Science [Internet]. 1998;281:1509 LP – 1512. Available from: http://science.sciencemag.org/content/281/5382/1509.abstract

54. Rafehi H, Orlowski C, Georgiadis GT, Ververis K, El-Osta A, Karagiannis TC. Clonogenic assay: Adherent cells. J Vis Exp. 2011;15–7.

55. Meeth K, Wang JX, Micevic G, Damsky W, Bosenberg MW. The YUMM lines: a series of congenic mouse melanoma cell lines with defined genetic alterations. Pigment Cell Melanoma Res. 2016;29:590–7.

56. Larue L. The WNT / Β - catenin pathway in melanoma. 2014;

57. Lucero OM, Dawson DW, Moon RT, Chien AJ. A Re-evaluation of the “Oncogenic” Nature of Wnt/β-catenin Signaling in Melanoma and Other Cancers. Curr Oncol Rep [Internet]. 2010;12:314–8. Available from: https://doi.org/10.1007/s11912-010-0114-3

58. Webster MR, Kugel CH, Weeraratna AT. The Wnts of change: How Wnts regulate phenotype switching in melanoma. Biochim Biophys Acta - Rev Cancer [Internet]. Elsevier B.V.; 2015;1856:244–51. Available from: http://dx.doi.org/10.1016/j.bbcan.2015.10.002

59. Kuphal S, Lodermeyer S, Bataille F, Schuierer M, Hoang BH, Bosserhoff AK. Expression of Dickkopf genes is strongly reduced in malignant melanoma. Oncogene. 2006;25:5027–36.

60. Mikheev AM, Mikheeva SA, Rostomily R, Zarbl H. Dickkopf-1 activates cell death in MDA-MB435 melanoma cells. Biochem Biophys Res Commun. 2007;352:675–80.

61. Haqq C, Nosrati M, Sudilovsky D, Crothers J, Khodabakhsh D, Pulliam BL, et al. The gene expression signatures of melanoma progression. Proc Natl Acad Sci USA. 2005;102:6092–7.

62. Gallagher SJ, Rambow F, Kumasaka M, Champeval D, Bellacosa A, Delmas V, et al. Beta-catenin inhibits melanocyte migration but induces melanoma metastasis. Oncogene [Internet]. 2013;32:2230–8. Available from: https://doi.org/10.1038/onc.2012.229

63. Widlund HR, Horstmann MA, Roydon Price E, Cui J, Lessnick SL, Wu M, et al. β-Catenin-induced melanoma growth requires the downstream target Microphthalmia-associated transcription factor. J Cell Biol. 2002;158:1079–87.

64. Hartman ML, Czyz M. MITF in melanoma: mechanisms behind its expression and activity. Cell Mol Life Sci [Internet]. 2014/11/30. Springer Basel; 2015;72:1249–60. Available from: https://pubmed.ncbi.nlm.nih.gov/25433395

65. Sierecki E, Sinko W, McCammon JA, Newton AC. Discovery of Small Molecule Inhibitors of the PH Domain Leucine-Rich Repeat Protein Phosphatase (PHLPP) by Chemical and Virtual Screening. J Med Chem [Internet]. American Chemical Society; 2010;53:6899–911. Available from: https://doi.org/10.1021/jm100331d

66. Mahadevan D, Powis G, Mash EA, George B, Gokhale VM, Zhang S, et al. Discovery of a novel class of AKT pleckstrin homology domain inhibitors. Mol Cancer Ther. 2008;7:2621–32.

67. Nawrotek A, Benabdi S, Niyomchon S, Kryszke MH, Ginestier C, Cañeque T, et al. PH-domain-binding inhibitors of nucleotide exchange factor BRAG2 disrupt Arf GTPase signaling. Nat Chem Biol [Internet]. Springer US; 2019;15:358–66. Available from: http://dx.doi.org/10.1038/s41589-019-0228-3

68. Chen Z, Wasney GA, Picaud S, Filippakopoulos P, Vedadi M, Angiolella VD, et al. Identification of a PGXPP degron motif in dishevelled and structural basis for its binding to the E3 ligase KLHL12. 2020;1–17.

69. Modell AE, Blosser SL, Arora PS. Systematic Targeting of Protein-Protein Interactions. Trends Pharmacol Sci [Internet]. 2016/06/04. 2016;37:702–13. Available from: https://pubmed.ncbi.nlm.nih.gov/27267699

70. Lai AC, Crews CM. Induced protein degradation: An emerging drug discovery paradigm. Nat Rev Drug Discov [Internet]. Nature Publishing Group; 2017;16:101–14. Available from: http://dx.doi.org/10.1038/nrd.2016.211

71. Verma R, Mohl D, Deshaies RJ. Harnessing the Power of Proteolysis for Targeted Protein Inactivation. Mol Cell [Internet]. Elsevier Inc.; 2020;77:446–60. Available from: https://doi.org/10.1016/j.molcel.2020.01.010

72. Simons M, Mlodzik M. Planar Cell Polarity Signaling: From Fly Development to Human Disease. Annu Rev Genet [Internet]. 2008;42:517–40. Available from: http://www.annualreviews.org/doi/10.1146/annurev.genet.42.110807.091432

73. Fellmann C, Hoffmann T, Sridhar V, Hopfgartner B, Muhar M, Roth M, et al. An optimized microRNA backbone for effective single-copy RNAi. Cell Rep. 2013;5:1704–13.

74. Rodgers L. Measurement of DNA Content Using Propidium Iodide (PI) Staining of Fixed Whole Cells. Cold Spring Harb Protoc [Internet]. 2006;2006:pdb.prot4436. Available from: http://cshprotocols.cshlp.org/content/2006/1/pdb.prot4436.short

