## Supplementary Information for "PLEKHA4 Promotes Wnt/β-catenin Signaling-Mediated G1/S Transition and Proliferation in Melanoma"

**Contents:**

Figures S1–S5
Table S1

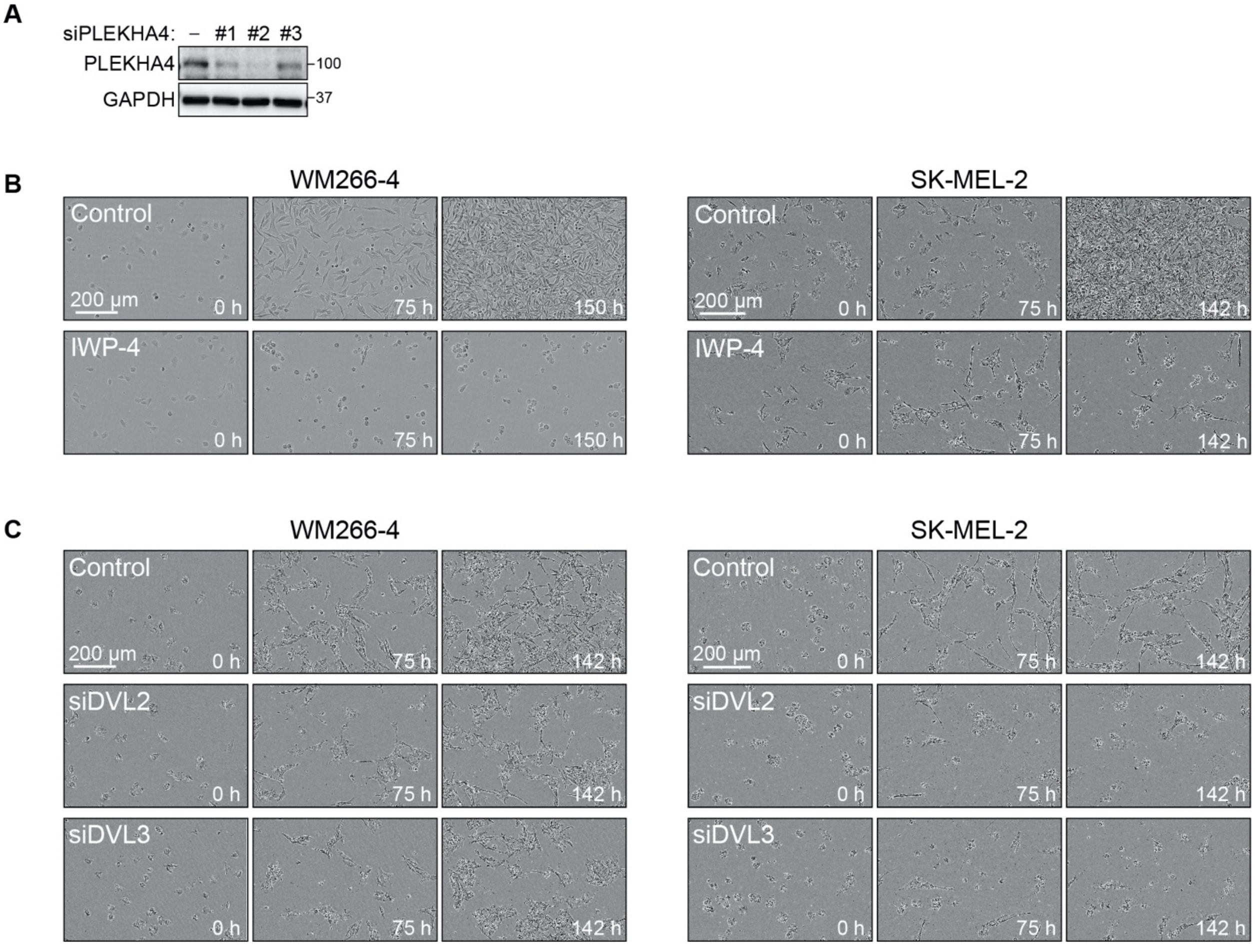


**Figure S1. Inhibition of Wnt signaling decreases proliferation in melanoma cells.** (A) Western blot validation of siRNA duplexes targeting different regions of PLEKHA4 (siPLEKHA4 #1, #2 and #3) or a negative control siRNA in WM266-4 melanoma cells. (B and C) Shown are representative brightfield images at the indicated timepoints of the IncuCyte proliferation assay of WM266-4 and SK-MEL-2 cells treated with the pan Wnt inhibitor IWP-4 or DMSO control (B) or with an siRNA duplex targeting DVL2 (siDVL2), DVL3 (siDVL3), or a negative control siRNA (C).

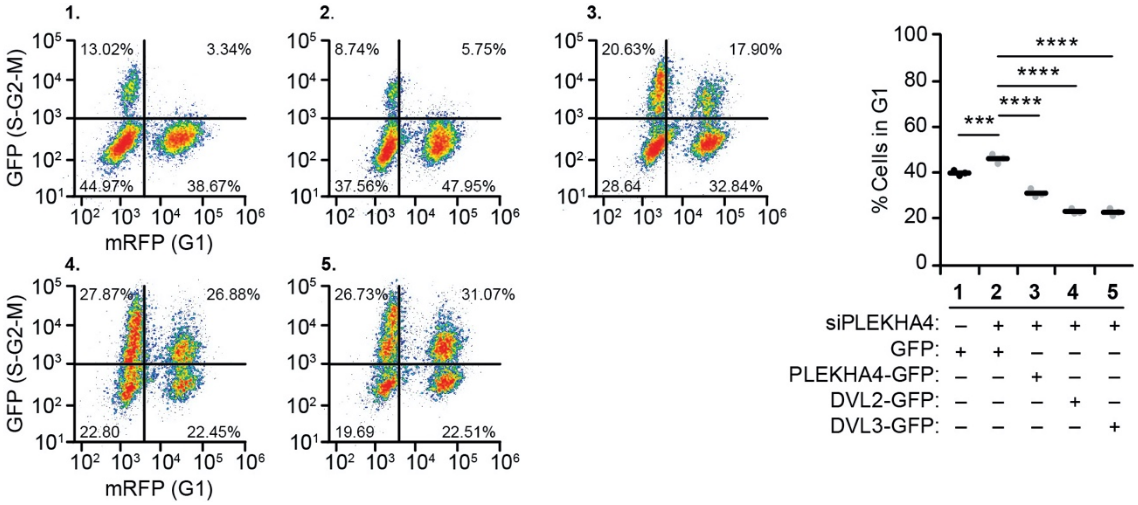


**Figure S2. PLEKHA4-GFP, DVL2-GFP and DVL3-GFP can rescue the attenuation of G1/S transition defect induced by PLEKHA4 knockdown.** WM266-4 cells were synchronized to G1 phase, subjected to siPLEKHA4 or negative control siRNA (–), and stimulated with media containing FBS and simultaneously transduced with conditioned media containing lentivirus encoding GFP, siRNA-resistant PLEKHA4-GFP, DVL2-GFP, or DVL3-GFP, followed by flow cytometry analysis. The plots at left show populations of cells expressing mRFP (G1) and GFP (S-G2-M), and the fraction of mRFP+ cells is plotted at right (n=3). *** $p < 0.001$, **** $p < 0.0001$.

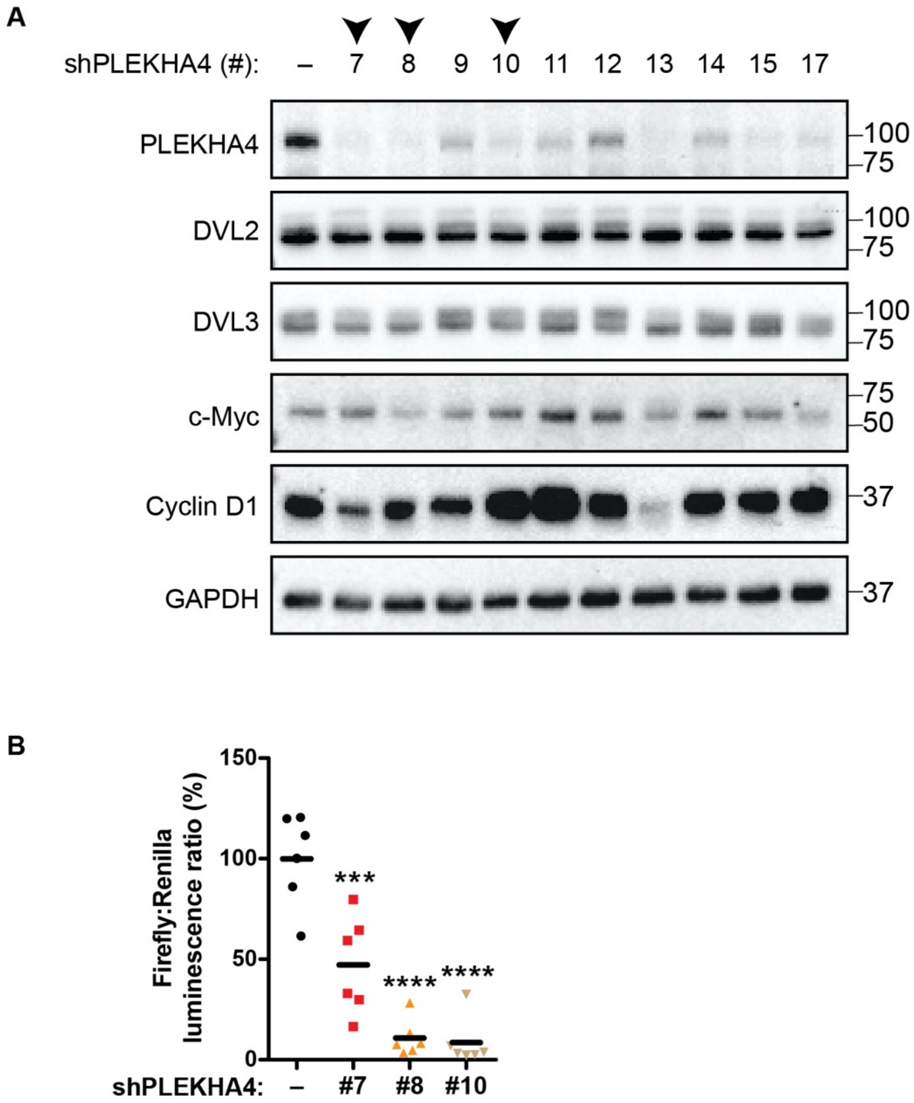


**Figure S3. Determination of optimal inducible shRNA constructs targeting human PLEKHA4.** WM266-4 cell lines were generated that stably expressed one of several different doxycycline-inducible shRNA constructs targeting human PLEKHA4. (A) Shown are representative Western blots of several WM266-4 shPLEKHA4 cell lines, evaluating levels of PLEKHA4 as well as Wnt signaling-related genes DVL2, DVL3, c-Myc, and Cyclin D1. Arrowheads indicate shRNAs #7, #8, and #10, which exhibited substantial decreases in levels of PLEKHA4 as well as DVL2, DVL3, Cyclin D1, and c-Myc in these cell lines, and were used for subsequent studies. (B) Wnt3a-stimulated β-catenin-dependent TOPFlash assay of the indicated WM266-4 cell lines expressing the indicated shPLEKHA4 or control shRNA construct (n=6). *** $p < 0.001$, **** $p < 0.0001$.

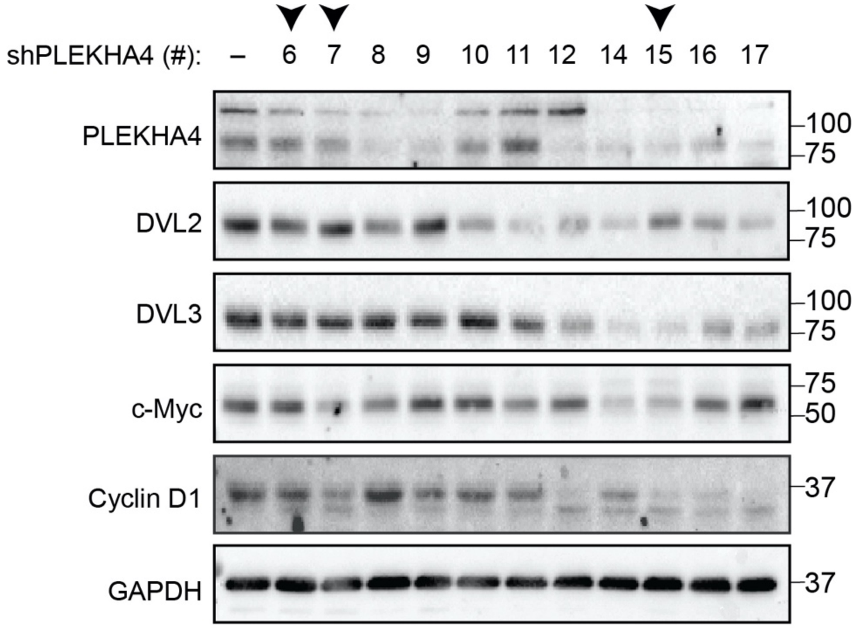


**Figure S4. Determination of optimal inducible shRNA constructs targeting mouse PLEKHA4.** YUMM1.7 cell lines were generated that stably expressed one of several different doxycycline-inducible shRNA constructs targeting mouse PLEKHA4. (A) Shown are representative Western blots of several WM266-4 shPLEKHA4 cell lines, evaluating levels of PLEKHA4 as well as Wnt signaling-related genes DVL2, DVL3, c-Myc, and Cyclin D1. Arrowheads indicate shRNAs #6, #7, and #15, which exhibited substantial decreases in levels of PLEKHA4 as well as DVL2, DVL3, Cyclin D1 and c-Myc in these cell lines and were used for subsequent studies.

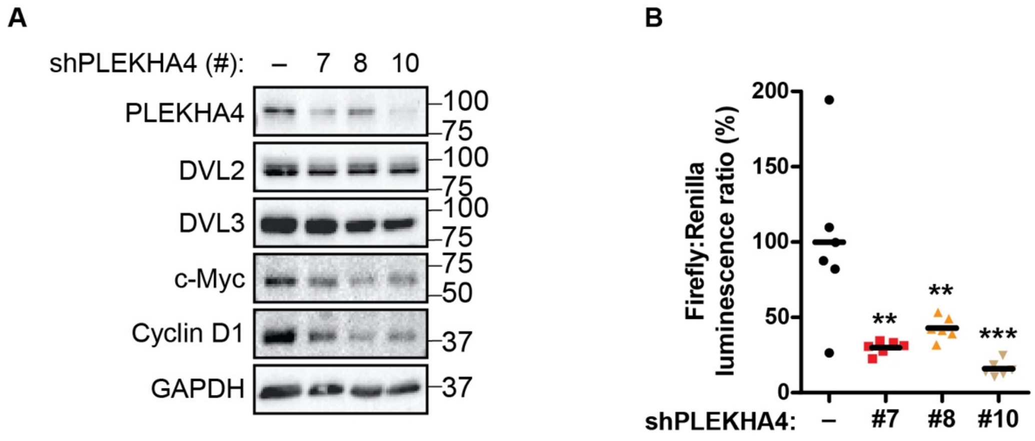


**Figure S5. Establishment of SK-MEL-2 cell lines stably expressing doxycycline-inducible shRNA against human PLEKHA4.** (A) Shown are representative Western blots of SK-MEL-2 cell lines stably expressing shPLEKHA4 #7, #8, and #10, evaluating levels of PLEKHA4 as well as Wnt signaling-related genes DVL2, DVL3, c-Myc, and Cyclin D1. Note that shRNAs #7, #8, and #10 exhibited decreases in levels of PLEKHA4 as well as DVL2, DVL3, Cyclin D1 and c-Myc in these cell lines. (B) Wnt3a-stimulated β-catenin-dependent TOPFlash assay of the indicated SK-MEL-2 cell lines expressing the indicated shPLEKHA4 or control shRNA construct (n=6). ** $p < 0.01$, ***, $p < 0.001$

**Table S1.** Sequences for siRNA[1] and shRNA[2].

| Name | Sequence |
| --- | --- |
| Negative control siRNA | Sense: rCrGrUrUrArArUrCrGrCrGrUrArUrArArUrArCrGrCrGrUAT |
| | Antisense: rArUrArCrGrCrGrUrArUrUrArUrArCrGrCrGrArUrUrArArCrGrArC |
| siPLEKHA4 #1 | Sense: rGrArArUrGrArGrArCrArGrArGrArCrUrUrArArGrGrArAGA |
| | Antisense: rUrCrUrUrCrCrUrUrArArGrUrCrUrCrUrGrUrCrUrCrArUrUrCrUrC |
| siPLEKHA4 #2 | Sense: rArGrCrUrArCrArArUrArUrUrArGrArCrCrArGrArUrGrGGC |
| | Antisense: rGrCrCrCrArUrCrUrGrGrUrCrUrArArUrArUrUrGrUrArGrCrUrGrG |
| siPLEKHA4 #3 | Sense: rUrCrUrCrArArCrArCrUrGrUrCrUrArArArUrUrUrGrGrATT |
| | Antisense: rArArUrCrCrArArArUrUrUrArGrArCrArGrUrGrUrUrGrArGrArArA |
| siDVL2 | Sense: rGrUrCrArCrGrCrUrArArArCrArUrGrGrArGrArArGrUrACA |
| | Antisense: rUrGrUrArCrUrUrCrUrCrCrArUrGrUrUrUrArGrCrGrUrGrArCrUrG |
| siDVL3 | Sense: rGrArUrArUrGrUrUrGrUrUrArCrArGrGrUrArArArCrGrAGA |
| | Antisense: rUrCrUrCrGrUrUrUrArCrCrUrGrUrArArCrArArCrArUrArUrCrUrC |
| Control shRNA (Renilla) | TGCTGTTGACAGTGAGCGCAGGAATTATAATGCTTATCTATAGTGAAGCCACAGATGTATAGATAAGCATTATAATTCCTATGCCTACTGCCTCGGA |
| Human shPLEKHA4 #7 | TGCTGTTGACAGTGAGCGCAGAGTCAACTTTCCACCAAAATAGTGAAGCCACAGATGTATTTTGGTGGAAAGTTGACTCTATGCCTACTGCCTCGGA |
| Human shPLEKHA4 #8 | TGCTGTTGACAGTGAGCGCAGCTACAATATTAGACCAGAATAGTGAAGCCACAGATGTATTCTGGTCTAATATTGTAGCTATGCCTACTGCCTCGGA |
| Human shPLEKHA4 #9 | TGCTGTTGACAGTGAGCGCACAGCTACAATATTAGACCAATAGTGAAGCCACAGATGTATTGGTCTAATATTGTAGCTGTATGCCTACTGCCTCGGA |
| Human shPLEKHA4 #10 | TGCTGTTGACAGTGAGCGCAGGTTCTCAGCCTCTCCCAAATAGTGAAGCCACAGATGTATTTGGGAGAGGCTGAGAACCTATGCCTACTGCCTCGGA |
| Human shPLEKHA4 #11 | TGCTGTTGACAGTGAGCGCACCGCGAGGAGAGTGTCCTAATAGTGAAGCCACAGATGTATTAGGACACTCTCCTCGCGGTATGCCTACTGCCTCGGA |
| Human shPLEKHA4 #12 | TGCTGTTGACAGTGAGCGCACAGCAGAGAGGAAAGAGAAATAGTGAAGCCACAGATGTATTTCTCTTTCCTCTCTGCTGTATGCCTACTGCCTCGGA |
| Human shPLEKHA4 #13 | TGCTGTTGACAGTGAGCGCAGCGAGTCACTCTGCTACAAATAGTGAAGCCACAGATGTATTTGTAGCAGAGTGACTCGCTATGCCTACTGCCTCGGA |
| Human shPLEKHA4 #14 | TGCTGTTGACAGTGAGCGAACAGATACGCTGCTGACCAAGTAGTGAAGCCACAGATGTACTTGGTCAGCAGCGTATCTGTCTGCCTACTGCCTCGGA |
| Human shPLEKHA4 #15 | TGCTGTTGACAGTGAGCGCAAGGAGGAGATAGACCAGAAATAGTGAAGCCACAGATGTATTTCTGGTCTATCTCCTCCTTATGCCTACTGCCTCGGA |
| Human shPLEKHA4 #17 | TGCTGTTGACAGTGAGCGCATCCACCATCTCCTCGCTCAATAGTGAAGCCACAGATGTATTGAGCGAGGAGATGGTGGATATGCCTACTGCCTCGGA |
| Mouse shPLEKHA4 #6 | TGCTGTTGACAGTGAGCGCAGCGCATGCGTAGAAACCAAATAGTGAAGCCACAGATGTATTTGGTTTCTACGCATGCGCTATGCCTACTGCCTCGGA |

| | |
|---|---|
| Mouse shPLEKHA4 #7 | TGCTGTTGACAGTGAGCGCACAGTGGATCTGCAGACTGA ATAGTGAAGCCACAGATGTATTCAGTCTGCAGATCCACTG TATGCCTACTGCCTCGGA |
| Mouse shPLEKHA4 #8 | TGCTGTTGACAGTGAGCGCAGATTCACCTTCACAGCAGA ATAGTGAAGCCACAGATGTATTCTGCTGTGAAGGTGAATC TATGCCTACTGCCTCGGA |
| Mouse shPLEKHA4 #9 | TGCTGTTGACAGTGAGCGCAGTCGATCTCACGTAATTTC ATAGTGAAGCCACAGATGTATGAAATTACGTGAGATCGAC TATGCCTACTGCCTCGGA |
| Mouse shPLEKHA4 #10 | TGCTGTTGACAGTGAGCGCACGGACACCAGAGCAGAGAA ATAGTGAAGCCACAGATGTATTTCTCTGCTCTGGTGTCCG TATGCCTACTGCCTCGGA |
| Mouse shPLEKHA4 #11 | TGCTGTTGACAGTGAGCGCAGGCTACTTCTGCTACCACA ATAGTGAAGCCACAGATGTATTGTGGTAGCAGAAGTAGCC TATGCCTACTGCCTCGGA |
| Mouse shPLEKHA4 #12 | TGCTGTTGACAGTGAGCGCAGGAGAAGAGTCCTCAGAGA ATAGTGAAGCCACAGATGTATTCTCTGAGGACTCTTCTCC TATGCCTACTGCCTCGGA |
| Mouse shPLEKHA4 #14 | TGCTGTTGACAGTGAGCGCGCTCTGTGTCACCTGGCTCAG TAGTGAAGCCACAGATGTACTGAGCCAGGTGACACAGAGCA TGCCTACTGCCTCGGA |
| Mouse shPLEKHA4 #15 | TGCTGTTGACAGTGAGCGAACAGATACGTTGTTGACTAAG TAGTGAAGCCACAGATGTACTTAGTCAACAACGTATCTGTC TGCCTACTGCCTCGGA |
| Mouse shPLEKHA4 #16 | TGCTGTTGACAGTGAGCGATTCCACCGTCTCCTCTCTGAG TAGTGAAGCCACAGATGTACTCAGAGAGGAGACGGTGGAAG TGCCTACTGCCTCGGA |
| Mouse shPLEKHA4 #17 | TGCTGTTGACAGTGAGCGACCATTGCCTCTTCTACTATAA TAGTGAAGCCACAGATGTATTATAGTAGAAGAGGCAATGGC TGCCTACTGCCTCGGA |

[1] The siRNA sequences were obtained as DsiRNA duplexes from IDT.
[2] For shRNA, the indicated 97-mers were designed and cloned into LT3GEPIR as described in Fellmann C *et al., Cell Rep* (2013) *5*, 1704–13.
